# The stage-specific regulation imposed by Importinβ-1 on HIV-1 propagation and infectivity dynamics

**DOI:** 10.1101/2024.06.10.598175

**Authors:** Sriram Yandrapally, Satarupa Sarkar, Shalini Saxena, Venkateshwarlu Naik, Srikanth Rapole, Sharmistha Banerjee

## Abstract

Dynamics of intra-host molecular evolution of viruses depend on the complexities of the cellular environments through which they transit. HIV infects several cell types in an infected human host, especially during chronic stages, which impose differential regulations on HIV persistence, driving it to latency, rapid propagation, or abortive infection. We observed that HIV-1 emerging from different cell-types differ in their encapsidated protein cargo, and infectivity. We investigated the role of Importinβ-1, which is encapsidated in virus emerging from CD4+T lymphocytes, but not in viruses from astrocytes. We deciphered that Importinβ-1 is packaged via interactions with HIV-1 Gag/Capsid. The encapsidated Importinβ-1 assisted nuclear-entry of the viral core and enhanced the infectivity during pre-integration stages. Conversely, high levels of endogenous Importinβ-1, which was observed to be induced upon infection and inflammatory stimulations, such as, IFNg/LPS treatments, restricted LTR-driven viral transcription during post-integration. The regulatory impact of Importinβ-1 was verified using primary CD4+T lymphocytes, thereby validating a non-canonical and novel role of Importinβ-1 as a restriction factor for HIV-1. Using deletion mutants, we demonstrate that the N-terminal domain of Importinβ-1 regulated viral transcription via SP1, and NRE regions of LTR. We propose that sequestering of Importinβ-1 by packaging in emerging virions thereby reducing its antiviral impact, is an adaptive strategy of a pathogen, supporting the concepts of evolutionary conflicts between viruses and hosts.

## Introduction

Integration of reverse-transcribed HIV RNA (proviral DNA) into the host genome is an essential step for establishment of HIV infection. The classical pathway used by HIV to infect a cell is via CD4 receptor and CXCR4/CCR5 co-receptors, which it crosses the cell membrane to enter the cytoplasm [1]. Similarly, to cross the nuclear membrane, HIV exploits the cytosolic receptors (Importins) and nucleoporins for its import into the nucleus of infected cell. The imports of viral factors and several host regulatory proteins are majorly taken care of by two families of cytosolic proteins, Importin-β, and Importin-α. Importinβ family of proteins are nucleocytoplasmic transport receptors (NTRs) and include proteins that can import and export different cargos through nuclear pore complexes (NPC) in cells. The family consists of twenty proteins; amongst them ten proteins are importers and seven proteins are exporters, two proteins are bidirectional, and one protein is uncharacterized [2]. Importinα family of proteins consist of seven proteins (Importin-α1-α7) and these function as adaptor proteins that connect Importinβ to their respective cargoes [3]. Import majorly takes place by two pathways, classical pathway and non-classical pathway. In the classical pathway, the prototype receptors of Importinα and Importinβ forms a heterodimeric complex to initiate the nuclear import. Whereas in non-classical pathway Importinβ-1 alone is sufficient to import the cellular cargoes from cytoplasm to nucleus [4]. Previous findings reported that Importin-7 imports HIV genomic DNA to nucleus in non-dividing cells, although this protein is not essential for the HIV infection [5]. Similarly, recent findings reported that HIV Capsid interacts with Importin-α and that mediates the nuclear entry of HIV in non-dividing cells [6]. Besides these reports, there are limited evidences on the role of Importins on nuclear import of HIV core/pre integration complex (PIC).

After successful entry and integration, HIV starts producing the viral copies by using its own 5’ LTR as a promoter and host transcription machinery. The HIV 5’LTR contains the enhancers and repressors elements that maintains the balance of HIV transcription, which indeed decides when to be active and when to be inactive [7,8]. Several DNA binding proteins have been shown to activate the viral transcription for establishing the infection. Amongst, NF-κB, SP1, and viral protein Tat, are very well studied [8]. Interestingly HIV Tat has been reported to both activate viral transcription and induce latency by inactivating the HIV LTR [9]. Similarly, few host DNA binding proteins that act as viral restriction factor are known to inactivate HIV LTR and induce latency [10,11]. However, HIV has evolved strategies to neutralize the impact of almost all host restriction factors for both establishing and re-establishing infections in a cell [12]. It is evident that HIV has a broad host cell tropism and infect numerous cell types in the host, each cell type exhibiting different regulatory impact on HIV propagation [13]. The replication of HIV is regulated by various combination of cellular and viral processes that include integration sites in the host genome, expression of various negative factors and positive factors in the infected cells and so on. The heterogeneity of expression yields different outcomes in the infected cells [14]. Some of the cell types reported to support high viral propagation are immune cells, such as, macrophages and CD4+ T lymphocytes, while cells residing in brain, such as, microglia and astrocytes, do not permit efficient viral propagation [13,15]. The phenotypic differences in propagation helps the virus in escaping the host selection pressure. While there are many studies on understanding the impact of cellular environment for viral replication [16–19] but infectivity dynamics and replication potential of the virus egressing from different producer cells are largely unknown. It is important to understand how transit through various cell types influences virus phenotype and infectivity, which may be navigating the latency or progression towards AIDS during chronic stages of infection.

In this study, we compared the infectivity dynamics of the virus emerging from CD4+T lymphocytes, a cell type supporting high HIV propagation, and astrocytes, a cell type that allow limited HIV propagation and attempted to understand the molecular basis of the differential infectivity. A mass spectrometry based proteomic analysis of HIV-1 emerging from CD4+ T lymphocytes and astrocytes revealed that the viruses differ in their encapsidated protein cargo. To validate that indeed the encapsidated host factors in emerging viruses make impact in subsequent infections and to establish a molecular link, we explored the role of Importin subunit beta-1 (here onward Importinβ-1 or Impβ-1), which was encapsidated in HIV-1 emerging from CD4+T lymphocytes, but not those from astrocytes. We show that the packaged Importinβ-1 enhanced viral infectivity during pre-integration stages by increasing the import of viral core to the nucleus, while post-integration, Importinβ-1 limited HIV-1 LTR activity, thereby reducing HIV-1 replication. The regulatory impact of Importinβ-1 was verified using primary CD4+T lymphocytes from four healthy donors. We also show that the regulatory impact imposed by Importinβ-1 is specific to HIV-1 LTR as SIV and SV40 virus promoters remained unaffected. In addition, we also present the domains of Importinβ-1 that are responsible for the regulatory impact on HIV LTR and the occupancy sites of Importinβ-1 on HIV LTR is SP1 and NRE sites. The study is the first report showing a stage-specific regulation imposed by a host factor on HIV-1 propagation. We propose a model which supports the notion of sequestering of Importinβ-1 by packaging in emerging virions as a survival strategy adapted by HIV.

## Results

### Virus egressing from different producer cells vary in their infectivity

Our previous observations pointed to differential spatial distribution of viral factors and released virus titers in CD4+T lymphocytes (SUP-T1) vs astrocytes (1321N1) [17]. We and others have shown that these two cell types, although are infected by HIV-1, have different regulation on HIV-1 propagation where CD4+T lymphocytes permitted high propagation of HIV-1, and astrocytes allowed limited propagation [16,20]. We also demonstrated that fundamental mechanisms, such as, viral RNA splicing is differentially regulated by HIV-1 Tat in these two cell types [16]. With these previous observations, we further investigated the phenotype and infectivity of HIV-1 population emerging from these two cell types. NL4.3 viruses were made by transfecting HEK293T cells with the plasmid pNL4.3. Virus was collected after 48 hours post-transfection. We refer to this as HEK293T_NL4.3. HEK293T_NL4.3 was then used to infect SUP-T1 and 1321N1 cell lines and the virus emerging from these cell types are referred to as SupT1_NL4.3, and 1321N1_NL4.3 respectively, here onwards. The infectivity of the viruses HEK293T_NL4.3, SupT1_NL4.3, and 1321N1_NL4.3 were then evaluated in TZM-bl cells using β-gal assays, in a concentration and time dependent manner. For viral infection, adherent TZM-bl cells were seeded 24 hours before infection and infected with 6ng/ml to 100ng/ml of p24 equivalent of HEK293T_NL4.3, SupT1_NL4.3, and 1321N1_NL4.3. The infectivity was tested by Tat-driven LTR-based β-galactosidase activity (Figure S1 A-C). It was observed that viruses generated from different cell types had different infectivity, which also showed a concentration dependence during infection. Comparatively, HEK293T_NL4.3 were highly infective in a dose dependent manner when assayed at 12 hours (Figure S1A), but it declined when assays were performed at later time points (Figure S1B, C). This may also reflect cell death due to high infectivity of HEK293T_NL4.3 at later time point which affected β-galactosidase readouts. A dose-dependent increase in the infectivity for SupT1_NL4.3 was observed at 24 hours, which we observed at 48 hours for 1321N1_NL4.3. As expected, a decreasing trend was observed for SupT1_NL4.3 and HEK293T_NL4.3 at 48hrs owing to cell death due to their higher infectivity. This overall suggested differential infectivity of HIV-1 derived from different cell types.

While viruses from different cell types were found to have different infectivity, it was possible that the same was due to the presence of cell-derived exosomes or secreted factors present in the virus preparations. To check the same, TZM-bl cells were infected with HEK293T_NL4.3 along with the uninfected SUP-T1 and 1321N1 culture supernatants (mock) (Figure S1D). 12 hours post-infection β-galactosidase assays were performed for infected TZM-bl cells. There were no significant differences in the infectivity of HEK293T_NL4.3 in the presence or absence of uninfected culture supernatants from either SUP-T1 or 1321N1 cells when compared with those infected with HEK293T_NL4.3 in the absence of any culture supernatants. The same was repeated with the SupT1_NL4.3 virus and 1321N1_NL4.3 viruses with exchange of the uninfected mock supernatants (Figure S1E, F).

With these observations, we concluded that the property of differential infectivity is associated with the virus enriched fractions, rather than culture supernatant. We speculated that these may be due to specific host factors that get associated with virus particles during egressing from different producer cells.

### Importinβ-1, a host factor found in HEK293T_NL4.3 and SupT1_NL4.3, but not in 1321N1_NL4.3 fractions, packages inside the virus by binding to the HIV-1 Gag/Capsid proteins

We next evaluated the host proteins that were enriched in HEK293T_NL4.3, SupT1_NL4.3, and 1321N1_NL4.3 fractions by mass spectrometry. The mass spectrometry based proteomic analysis indicated differential host proteins associated with these viruses, some of the significantly represented proteins are listed Table (Table S1, Supplementary File 2). Some of the proteins identified as host cargo in emerging virus population are already reported to influence HIV-1 infection cycle [21]. In this study, we focused on Importinβ-1, a host protein associated with the HEK293T_NL4.3 and SupT1_NL4.3 but not with the 1321N1_NL4.3. The mass spectrometry data was re-confirmed by western blot using anti-Importinβ-1 antibodies (Figure 1A-C).

**Figure 1:**
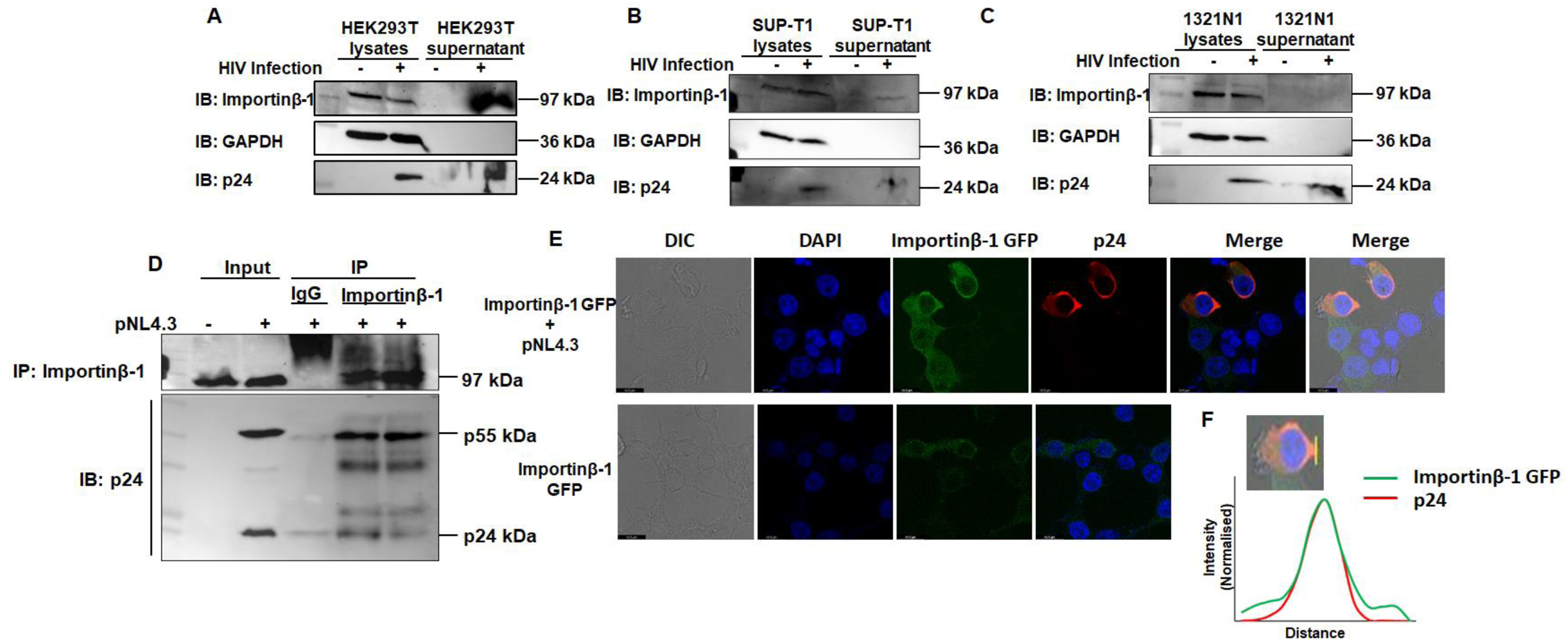
Encapsidation of Importinβ-1 in emerging viruses. **(A)** Immunoblot showing the presence of Importinβ-1 in virus preparation from culture supernatants of pNL4.3 transfected HEK293T cells. The presence of virus was detected with anti-p24 antibody. GAPDH was used as the internal control. (**B)** Immunoblot showing the presence of Importinβ-1 in virus preparation from culture supernatants of NL4.3 infected SUP-T1 cells. The presence of virus was detected with anti-p24 antibody. GAPDH was used as the internal control. **(C)** Immunoblot showing the presence of Importinβ-1 in virus preparation from culture supernatants of NL4.3 infected 1321N1 cells. The presence of virus was detected with anti-p24 antibody. GAPDH was used as the internal control. **(D)** Co-immune precipitation showing Importinβ-1 interaction with Gag and Capsid proteins upon pNL4.3 transfection in HEK293T cells. Mouse IgG was used as an isotypic control. **(E)** Confocal microscopy showing co-localisation of Importinβ-1 GFP and Gag/Capsid protein in the HEK293T cells. **(F)** The plot profiles showing the overlapping of Importinβ-1 GFP and Gag/Capsid protein in representative single cell generated using ImageJ software. All experiments were done at least 3 times.

Understanding that a host protein will have a higher chance of getting encapsidated in emerging virus if it interacted with viral factors, the interactions of Importinβ-1with HIV-1 Gag-Pol or Capsid proteins were checked by co-immunoprecipitation with either endogenous Importinβ-1 or HIV-1 Capsid from pNL4.3 transfected HEK293T cells. We observed that Importinβ-1 indeed interacted with Gag (p55) and Capsid (p24) of HIV-1 (Figure 1D, Figure S2A). These associations were also verified microscopically upon transiently expressing Importinβ-1 in pNL4.3 transfected HEK293T cells, where Importinβ-1-GFP colocalized with Capsid (p24) (Figure 1E, F). An interesting observation was higher intensity, and higher localization, of Importinβ-1-GFP at the nuclear envelope in the infected cells compared to uninfected cells (Figure 1E). Similarly, we have checked endogenous Importinβ-1 localisation with and without infection (Figure S2B)

Once confirmed that Importinβ-1 is indeed associated with HEK293T_NL4.3 and SupT1_NL4.3, and is packaged into these virions by virtue of its interactions with Gag and Capsid, we sought to investigate that why Importinβ-1 is not packaged in 1321N1_NL4.3. Towards this, we checked the levels of the Importinβ-1 in the HEK293T, SUP-T1, 1321N1 cells with and without HIV-1 infections. We observed that Importinβ-1 levels were much lower in 1321N1 cells as compared to HEK293T, and SUP-T1 (Figure S2C, D).

With this we concluded that Importinβ-1 packages inside the viruses by binding to the Gag / Capsid protein of HIV-1, and the absence of Importinβ-1 in 1321N1_NL4.3 may be because of its lower expression in 1321N1.

### Packaged Importinβ-1 enhances viral infectivity by augmenting the viral core transport into the nucleus of infected host cells

Next, we investigated the impact of the packaged Importinβ-1 in HEK293T_NL4.3 and SupT1_NL4.3. As it is known that the principal function of Importinβ-1 is to import proteins from cytoplasm to nucleus, we studied the virus import in the infected cells from cytoplasm to nucleus by measuring the presence of two LTR circles, Integrase, and Capsid in the nucleus of the TZM-bl cells infected with either HEK293T_NL4.3, SupT1_NL4.3 or 1321N1_NL4.3 at 8 or 12 hrs post-infection. Microscopic visualization using immunofluorescence showed that Capsid protein levels were much higher in the nucleus of TZM-bl cells infected with HEK293T_NL4.3 and SupT1_NL4.3 than the 1321N1_NL4.3 after 8 hrs of infection (Figure S3A). The ratio of nuclear to cytoplasmic localized Capsid was then quantified in TZM-bl cells infected with either HEK293T_NL4.3, SupT1_NL4.3 or 1321N1_NL4.3, which confirmed that proportionately, Capsids entered more into the nucleus of HEK293T_NL4.3, and SupT1_NL4.3 infected cells (Figure S3A). Further, it was observed that two LTR circles were less in the 1321N1_NL4.3 infected TZM-bl cells compared to HEK293T_NL4.3 and SupT1_NL4.3 infected cells 12 hrs post-infection (Figure S3B). Similar observations were made for the levels of Integrase protein in the infected TZM-bl cells through nucleus and cytoplasm fractionation followed by western blot using anti-Integrase antibodies (Figure 2B). We found, like other parameters, HEK293T_NL4.3 and SupT1_NL4.3 infected cells showed higher levels of Integrase protein in the nucleus of infected TZM-bl cells than the 1321N1_NL4.3 infected cells (Figure 2B, Figure S3C). The western-blot quantifications of integrase protein in the nucleus and cytoplasm 8hrs post-infection are shown in Figure 2B.

**Figure 2:**
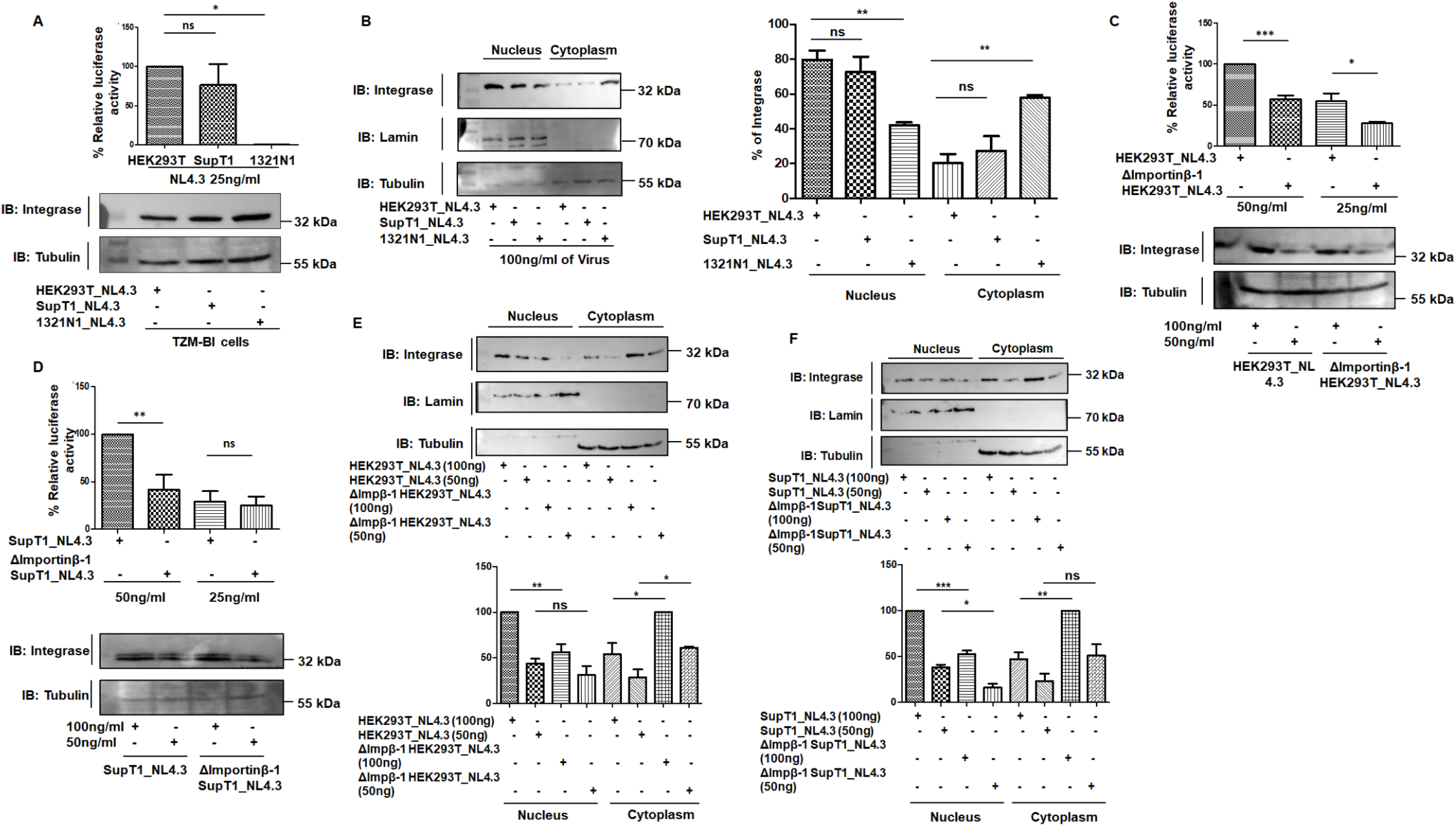
Encapsulated Importinβ-1 is critical for import of viral core to nucleus. **(A)** LTR-driven luciferase activity with the HEK293T_NL4.3, SupT1_NL4.3, 1321N1_NL4.3 viruses after the 16hrs of post infection in TZM-bl cells. The corresponding immunoblot showing the levels of Integrase. Tubulin was used as a loading control. **(B)** Representative immunoblot showing Integrase in the nuclear and cytoplasmic fractions of the infected TZM-bl cells, after 8 hrs of infection. Lamin and Tubulin were used as a nuclear and cytoplasmic controls respectively. Bar graph showing immunoblot quantification of Integrase in the nuclear and cytoplasmic fractions of three independent experiments. **(C)** LTR-driven luciferase activity with the HEK293T_NL4.3, ΔImportinβ-1 HEK293T_NL4.3, viruses after the 16hrs of post infection. The corresponding immunoblot showing the levels of Integrase. Tubulin was used as a loading control. **(D)** LTR-driven luciferase activity with the SupT1_NL4.3, ΔImportinβ-1 SupT1_NL4.3, viruses after the 24hrs of post infection. The corresponding immunoblot showing the levels of Integrase. Tubulin was used as a loading control. **(E)** Representative immunoblot showing the Integrase protein in the nucleus and cytoplasm of TZM-bl cells infected with either HEK293T_NL4.3 or ΔImpβ-1 HEK293T_NL4.3 viruses (100ng/ml or 50ng/ml), after 8 hrs of infection. Lamin and Tubulin were used as a nuclear and cytoplasmic controls respectively. Bar graph showing immunoblot quantification of Integrase in the nuclear and cytoplasmic fractions of three independent experiments. **(F)** Representative immunoblot showing the Integrase protein in the nucleus and cytoplasm of TZM-bl cells infected with either SupT1_NL4.3 or ΔImpβ-1 SupT1_NL4.3 viruses (100ng/ml or 50ng/ml), after 8 hrs of infection. Lamin and Tubulin were used as a nuclear and cytoplasmic controls respectively. Bar graph showing immunoblot quantification of Integrase in the nuclear and cytoplasmic fractions of three independent experiments. All experiments were done at least 3 times. The significance is determined using an unpaired student’s t-test. The p values are denoted as *** p <0.001; ** p <0.01; * p <0.05, while non-significant values are denoted by n.s.

The above finding suggested that Importinβ-1 packaging had significant impact on the viral core import from cytoplasm to nucleus at an early stage of HIV-1 lifecycle. To further confirm the same, we made viruses deficient in packaged Importinβ-1 from HEK293T and SUP-T1 cells, by knocking out Importinβ-1 from these cells (Figure S3D). These virions are referred to as ΔImportinβ-1 HEK293T_NL4.3 and ΔImportinβ-1 SupT1_NL4.3 viruses in the following experiments. The infectivity of these Importinβ-1 deficient viruses was then compared with HEK293T_NL4.3 and SupT1_NL4.3 in TZM-bl cells, using the same three parameters as detailed above. The results clearly showed that, when compared to HEK293T_NL4.3 and SupT1_NL4.3 infections, ΔImportinβ-1-HEK293T_NL4.3, and ΔImportinβ-1-SupT1_NL4.3 viruses had reduced infectivity (Figure 2C, D), reduced Integrase in the nucleus of infected cells (Figure 2E, F), reduced Capsid levels in the nucleus of infected cells (Figure S3E), and reduced two LTR circles (Figure S3F).

With these experiments, we inferred that as HEK293T_NL4.3 and SupT1_NL4.3 viruses carried Importinβ-1, more viral cores could be imported into the nucleus of the infected TZM-bl cells, thereby increasing the overall infectivity of these viruses. Concomitantly, the same events were not noticed in the viruses deficient in encapsidated Importinβ-1, including 1321N1_NL4.3, ΔImportinβ-1-HEK293T_NL4.3, and ΔImportinβ-1-SupT1_NL4.3, where viral cores were seen lesser in the nucleus, thus explaining their reduced infectivity.

### Viral encapsidated Importinβ-1 compensates for low endogenous Importin**β**-1 in transporting viral core into the nucleus of the infected cells during early stages of the viral lifecycle

Since our findings revealed that encapsidated Importinβ-1 helped viral core import, we enquired if this will be an advantage for the virus in a cell expressing lower levels of Importinβ-1. To mimic the same, we generated knockdown (KD) of Importinβ-1 in the TZM-bl cells (Figure S4A). Then wild-type TZM-bl and KD TZM-bl cells were infected with the Importinβ-1 containing viruses, HEK293T_NL4.3, and SupT1_NL4.3, and Importinβ-1 deficient viruses 1321N1_NL4.3, ΔImportinβ-1 HEK293T_NL4.3, and ΔImportinβ-1 SupT1_NL4.3. The viral core entry into the nucleus of infected cells were then scored by quantifying proportion of integrase and Capsid in the nucleus after 8hrs of infection and infectivity of the viruses were scored by Tat-dependent LTR-driven luciferase activity. Our fractionation studies showed that there is a reduction in the intra-nuclear integrase levels in Importinβ-1 KD TZM-bl cells infected with HEK293T_NL4.3, SupT1_NL4.3 by 32%, and 26% (Figure 3A,B), respectively, but for 1321N1_NL4.3, ΔImportinβ-1 HEK293T_NL4.3, and ΔImportinβ-1 SupT1_NL4.3, the reduction was much higher amounting to 65%, 72% and 71% respectively (Figure 3C-E) when compared with that from infected wild-type TZM-bl cells. Similarly, Capsid proteins in the nucleus of these infected cells showed the same trend as integrase, with highly reduced Capsids in the nucleus of the cell deficient in endogenous Importinβ-1 and infected with viruses deficient in encapsidated Importinβ-1 (Figure S4B-D). Supporting these, the Tat-dependent LTR-driven luciferase activity showed lowest infectivity when there is low Importinβ-1 in cells and no Importinβ-1 in viruses (Figure S5A-E).

**Figure 3:**
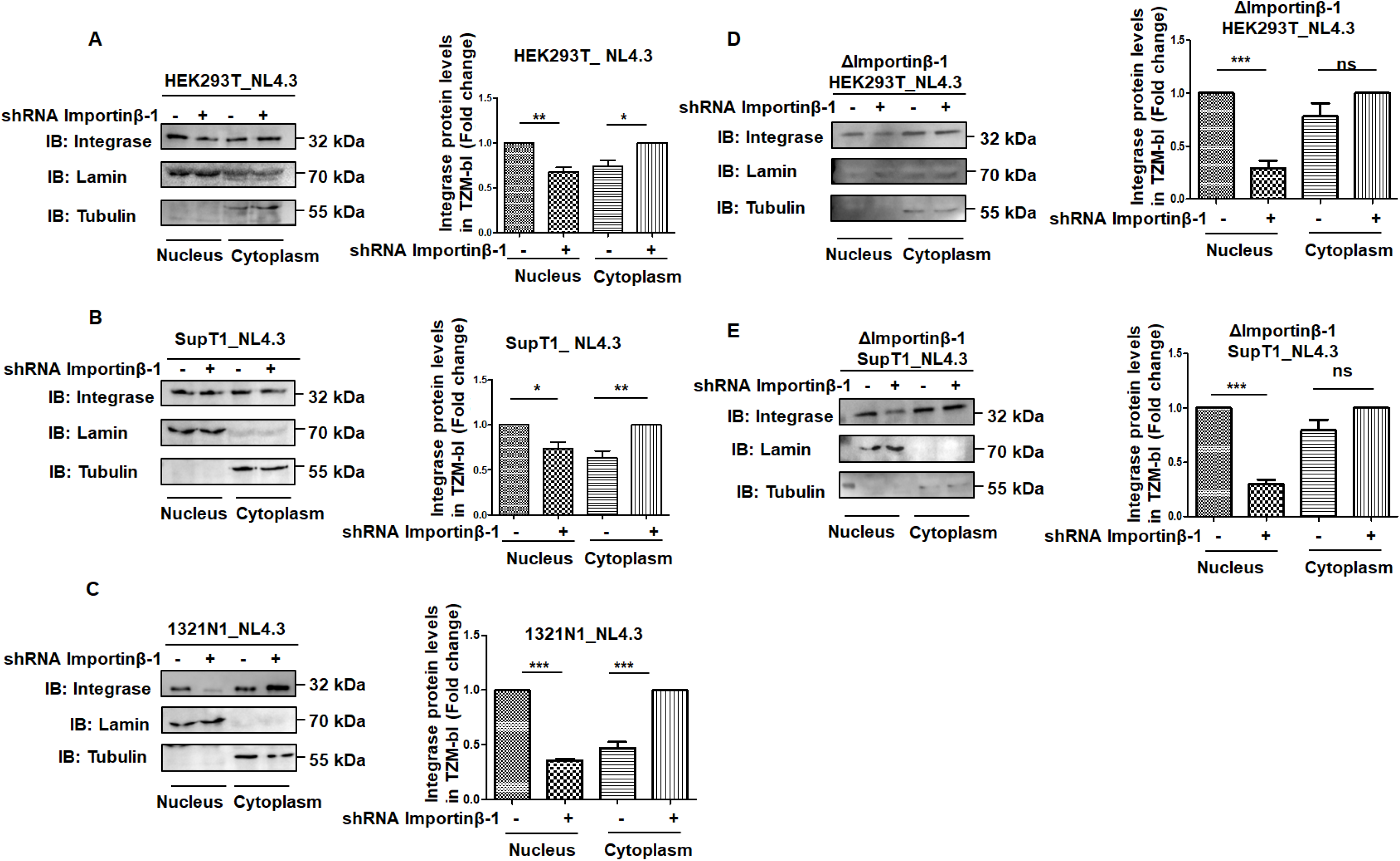
Endogenous or virus encapsidated Importinβ-1 support import of viral core to the nucleus of infected cells. **(A)** Representative immunoblot showing Integrase in the nuclear and cytoplasmic fractions of control shRNA or Importinβ-1 shRNA treated TZM-bl cells infected with HEK293T_NL4.3 viruses, after 8hrs of infection. Lamin and Tubulin were used as a nuclear and cytoplasmic controls respectively. Bar graph showing immunoblot quantification of Integrase in the nuclear and cytoplasmic fractions of three independent experiments. **(B)** Representative immunoblot showing Integrase in the nuclear and cytoplasmic fractions of control shRNA or Importinβ-1 shRNA treated TZM-bl cells infected with SupT1_NL4.3 viruses, after 8hrs of infection. Lamin and Tubulin were used as a nuclear and cytoplasmic controls respectively. Bar graph showing immunoblot quantification of Integrase in the nuclear and cytoplasmic fractions of three independent experiments. **(C)** Representative immunoblot showing Integrase in the nuclear and cytoplasmic fractions of control shRNA or Importinβ-1 shRNA treated TZM-bl cells infected with 1321N1_NL4.3 viruses, after 8hrs of infection. Lamin and Tubulin were used as a nuclear and cytoplasmic controls respectively. Bar graph showing immunoblot quantification of Integrase in the nuclear and cytoplasmic fractions of three independent experiments. **(D)** Representative immunoblot showing Integrase in the nuclear and cytoplasmic fractions of control shRNA or Importinβ-1 shRNA treated TZM-bl cells infected with ΔImportinβ-1 HEK293T_NL4.3 viruses, after 8hrs of infection. Lamin and Tubulin were used as a nuclear and cytoplasmic controls respectively. Bar graph showing immunoblot quantification of Integrase in the nuclear and cytoplasmic fractions of three independent experiments. **(E)** Representative immunoblot showing Integrase in the nuclear and cytoplasmic fractions of control shRNA or Importinβ-1 shRNA treated TZM-bl cells infected with ΔImportinβ-1 SupT1_NL4.3 viruses, after 8hrs of infection. Lamin and Tubulin were used as a nuclear and cytoplasmic controls respectively. Bar graph showing immunoblot quantification of Integrase in the nuclear and cytoplasmic fractions of three independent experiments. All experiments were done at least 3 times. The significance is determined using an unpaired student’s t-test. The p values are denoted as *** p <0.001; ** p <0.01; * p <0.05, while non-significant values are denoted by n.s.

Overall, these findings suggested that encapsidated Importinβ-1 helped HIV-1 infectivity by facilitating import of viral core even under the condition with compromised endogenous Importinβ-1 levels during early phase of viral lifecycle. Although HIV-1 also uses endogenous cellular Importinβ-1 when it does not carry it but the import of viral core into the nucleus is not as augmented as when it carries Importinβ-1.

### Intracellular Importin**β**-1 levels inversely correlated with released virions during infection

Having established the advantage of encapsidated Importinβ-1 in the early stages of HIV-1 lifecycle, we next focused on endogenous levels of Importinβ-1 during the course of infection. Expression of Importinβ-1 in SUP-T1 cells upon infection with NL4.3 or in HEK293T cells upon transfection with pNL4.3 vector was quantified in a time-dependent manner every 24 hrs from 0 hrs to until 96 hrs post infection/transfection. We observed that there is an increase in Importinβ-1 levels during the early stage of infection to till 24hrs post infection/transfection but it decreased continually till 96hrs post infection/transfection (Figure 4A, B, Figure S6A-C). The released virions, in terms of p24 equivalent, expectedly, continued to increase (Figure 4C, Figure S6C), pointing to an inverse relation between endogenous Importinβ-1 levels and released virions in both SUP-T1 and HEK293T cells (Figure 4D). With this, we inferred that Importinβ-1 levels were altered upon NL4.3 infection or pNL4.3 transfection.

**Figure 4:**
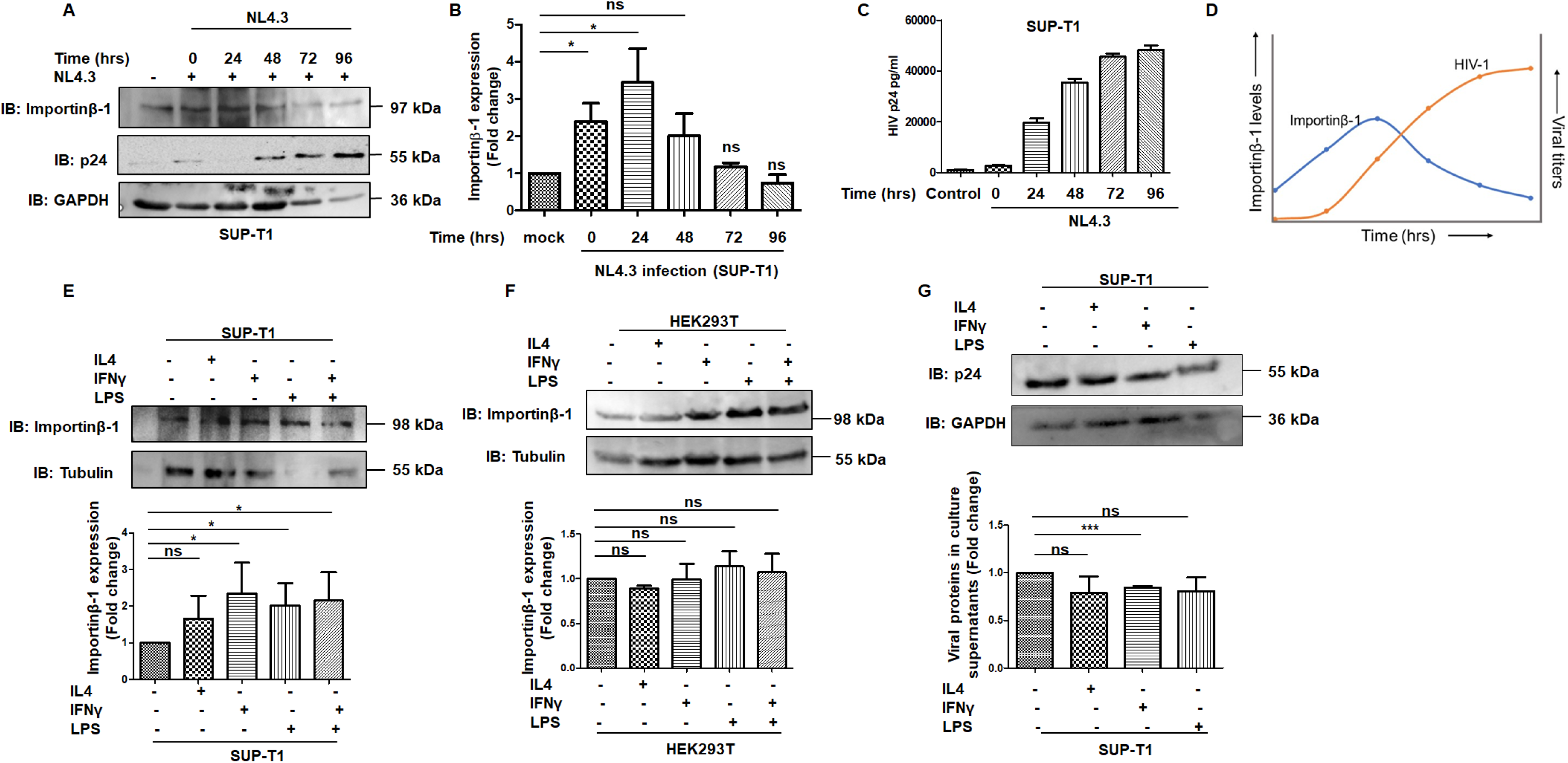
Converse levels of Importinβ-1 to HIV titers during infection or cytokine stimulation. **(A)** Representative Immunoblot showing the levels of Importinβ-1 from 0hrs to 96hrs in SUP-T1 cells upon NL4.3 infection. GAPDH was used as a control. **(B)** Bar graph representing the quantification of Importinβ-1 for the experiments represented in A. **(C)** Levels of released virus determined by p24 ELISA in the supernatant of NL4.3 infected SUP-T1 cells from 0 hrs to 96 hrs. **(D)** Line graph representing the relationship of endogenous Importinβ-1 and HIV titers in SUP-T1 cells during the course of HIV infection from 0 hrs to 96 hrs. Endogenous GAPDH was used as a loading control. **(E)** Representative Immunoblot showing the levels of Importinβ-1 in SUP-T1 cells upon treatment with IFNγ, LPS, and IL4. Tubulin was used as a control. Bar graph showing immunoblot quantification of Importinβ-1 from three independent experiments. **(F)** Representative Immunoblot showing the levels of Importinβ-1 in HEK293T cells upon treatment with IFNγ, LPS, and IL4. Tubulin was used as a control. Bar graph showing immunoblot quantification of Importinβ-1 from three independent experiments. **(G)** Representative Immunoblot showing the levels of p55 in SUP-T1 cells upon treatment with IFNγ, LPS, and IL4. GAPDH was used as a control. Bar graph showing immunoblot quantification of Importinβ-1 from three independent experiments. All experiments were done at least 3 times. The significance is determined using an unpaired student’s t-test. The p values are denoted as *** p <0.001; * p <0.05, while non-significant values are denoted by n.s.

We next checked if Importinβ-1 expression changes with inflammatory microenvironment. To study this, SUP-T1 cells and HEK293T cells were treated with IL-4 (40ng/ml), IFN-γ (20ng/ml), and LPS (2ug/ml) and Importinβ-1expression levels were analysed after 24 h to 30 h post treatment. We observed that Importinβ-1 expression was enhanced upon treatment with IFN-γ, LPS more as compared IL-4 and PBS in SUP-T1 cells but not in HEK293T (Figure 4E, F). With this, we inferred that Importinβ-1 expression levels in a cell increases when it faces an inflammatory microenvironment. Similarly, HIV replication was checked in IL-4, IFN-γ, and LPS treated SUP-T1, wherein we observed that HIV replication was reduced as compared to the control cells (Figure 4G).

This data was very intriguing as we observed an increase in Importinβ-1, both upon HIV infection and cytokine stimulations, correlated with decreased viral titers. This prompted us to investigate the impact of intracellular Importinβ-1 on HIV-1 lifecycle, focussing on post-integration stages.

### Importinβ-1 is a potential restriction factor for HIV-1 during post-integration stages in primary CD4+ T lymphocytes

To understand the role of Importinβ-1 on post-integration stages of HIV-1 lifecycle, we over-expressed Importinβ-1 in HEK293T cells and viral transcripts and protein levels were checked after 48 hrs. We observed that Importinβ-1 over-expression inhibited the replication of NL4.3, NL-ADA8 and IndieC but not SIV. In fact, Importinβ-1 over-expression had a positive impact on SIV replication (Figure 5A-C). Further, we evaluated the impact of dose-dependent increase in Importinβ-1 levels on intra-cellular p55 and the released p24, wherein, we observed a gradual decrease in the intracellular p55 and extracellular p24 levels (Figure 5D, E). To verify this, Importinβ-1 was both knocked down and knocked out in HEK293T cells after episomal expression of pNL4.3. We observed that Importinβ-1 knockdown resulted in elevated intracellular p55 and extracellular p24 levels but its knockout resulted in reduced p55 and extracellular p24 levels (Figure S7A, B). We reasoned that as Importinβ-1 aids in internalization of both viral proteins and host transcription factors into the nucleus of an infected cell that drive HIV-LTR [22,23], complete knockout might be detrimental to virus. Additionally, we observed a dose-dependent decrease in intracellular p55 and extracellular p24 levels with increased knocked-down levels of Importinβ-1 performed by gradually increasing the concentration of Importinβ-1-targeting shRNA in HEK293T cells (Figure S7C, D).

**Figure 5:**
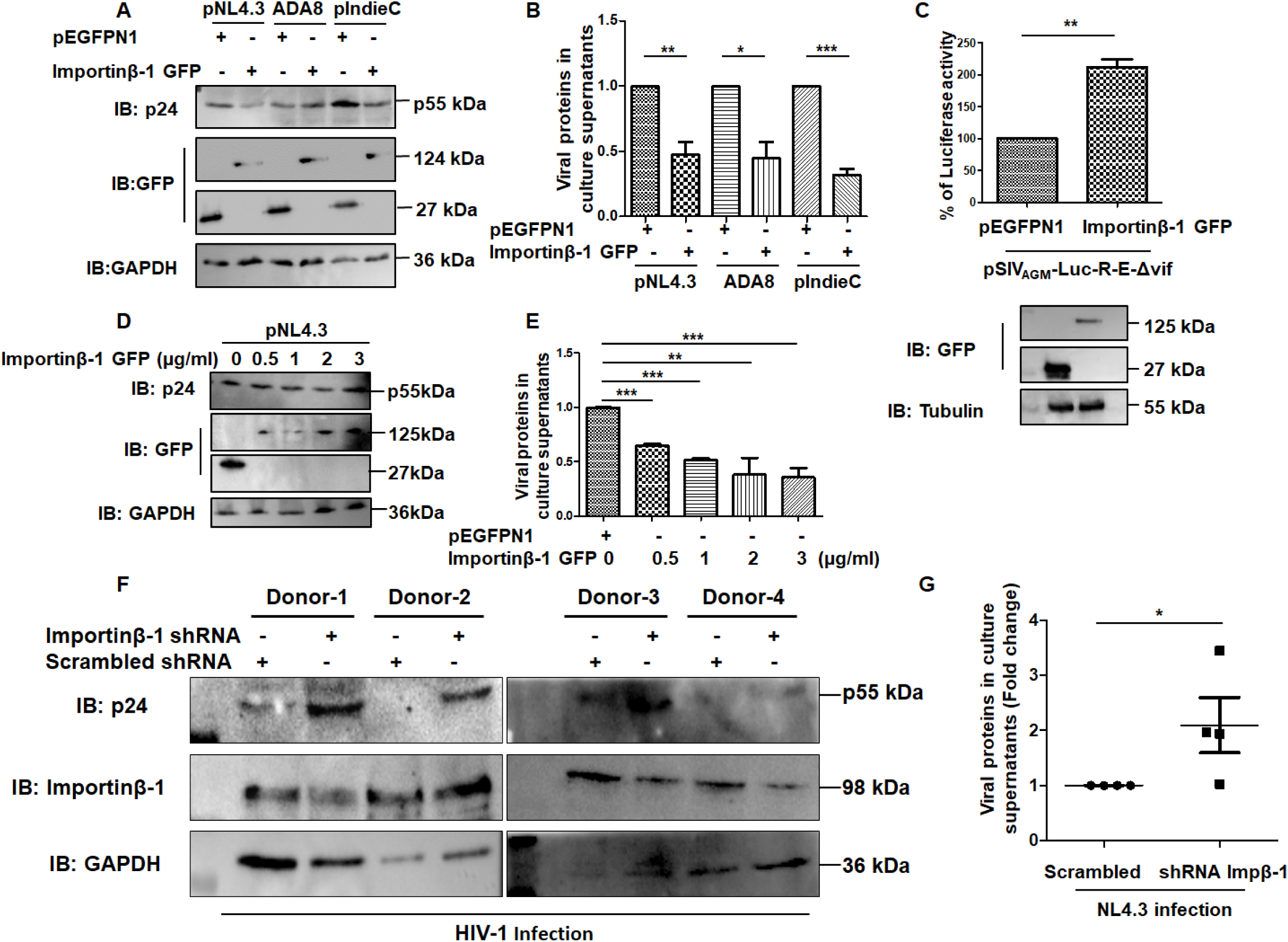
Importinβ-1 is a potential restriction factor of HIV. **(A)** Representative immunoblot showing the intracellular expression of p55 levels in HEK293T cells upon transiently expressing Importinβ-1 or respective vector control along with different viral constructs. **(B)** Levels of released virus determined by p24 ELISA in supernatant of HEK293T cells upon transiently expressing Importinβ-1 or respective vector control along with different viral constructs. **(C)** The bar graph representing the luciferase activity in HEK293T cells upon co-transfecting Importinβ-1 GFP or vector control with pSIVAGM-Luc-R-E-ΔVif. **(D)** Representative immunoblot showing the intracellular expression of p55 levels in HEK293T cells upon co-transfection of pNL4.3 with increasing concentration of Importinβ-1 GFP construct (0.5, 1, 2, 3 µg/ml). GAPDH was used as a loading control. **(E)** Levels of released virus determined by p24 ELISA in supernatant of pNL4.3 transfected HEK293T cells upon co-transfection of pNL4.3 with increasing concentration of Importinβ-1 GFP construct (0.5, 1, 2, 3 µg/ml). **(F)** Immunoblot showing the intracellular expression of p55 levels in HIV infected primary CD4+T lymphocytes upon Importinβ-1 specific shRNA transduced primary cells. **(G)** Levels of released virus determined by p24 ELISA in supernatant of HIV infected primary CD4+T lymphocytes upon Importinβ-1 specific shRNA transduction. All experiments were done at least 3 times. The significance is determined using an unpaired student’s t-test and Mann Whitney test was executed for figure G. The p values are denoted as *** p <0.001; ** p <0.01; * p <0.05, while non-significant values are denoted by n.s.

To substantiate these experiments in physiologically relevant conditions, we used primary CD4+T lymphocytes from four healthy donors. Primary CD4+T lymphocytes were first infected with NL4.3 virus and after 24hrs of infection, knockdown of Importin-β-1 was initiated. This experimental set-up allowed the presence of intracellular Importinβ-1 during pre-integration stages, but lowered Importinβ-1 post-integration of proviral DNA. Under these conditions, we observed that Importinβ-1 knockdown resulted in elevated intracellular p55 and extracellular p24 levels (Figure 5F, G). Importinβ-1 knockdown and knockout data pointed that some basal levels of Importinβ-1 is required for HIV replication and high Importinβ-1 levels negatively affected the viral titers.

### Importinβ-1 inhibits HIV replication by negatively regulating HIV-1 LTR activity via SP1 and NRE sites on LTR through its N and C-terminal domain

With Importinβ-1 functioning as a restriction factor for HIV replication in post-integration stages of life cycle, we further investigated the possible mechanism of Importinβ-1 mediated inhibition. As it is known that Importinβ-1 is involved in import of host factors from the cytoplasm to the nucleus, we initially checked if there is any perturbation in the export of HIV-1 RNA from the nucleus to the cytoplasm. We over expressed Importinβ-1 and respective vector controls along with pNL4.3 in HEK293T cells. After 48 h of transfection, the nuclear and cytoplasmic fractions were separated, followed by RNA isolation from these fractions. We then analysed the distribution of partially spliced and unspliced viral RNA in nuclear and cytoplasmic fraction through RRE-RNA qRT-PCR. We observed an increased RRE-RNA levels in the cytoplasm upon Importinβ-1 over-expression (Figure S8A), however the total viral RNA in Importinβ-1 over-expression background was reduced (Figure S8B). Thus, we could assess that the negative regulation on HIV-1 levels by Importinβ-1 is not mediated by affecting nucleocytoplasmic transport of viral RNA, perhaps it is regulated at the transcriptional level.

We then evaluated the HIV-1 LTR promoter strength upon ectopic expression of Importinβ-1 in HEK293T cells, in the presence or absence of Tat. This was performed by transfecting HEK293T cells with LTR-luc and Importinβ-1. Controls were used as required. In this experiment, we observed that Importinβ-1 expression reduced the LTR-driven luciferase activity, which was similar in the presence or absence of Tat (Figure 6A). However, we observed increase in luciferase activity upon increasing Tat concentration in the background of Importinβ-1 ectopic expression (Figure 6B). To verify if the impact of Importinβ-1 is restricted to HIV-1 LTR, we performed these assays on SV40 viral promoter. Importinβ-1 had no effect on SV40 promoter (Figure 6C).

**Figure 6:**
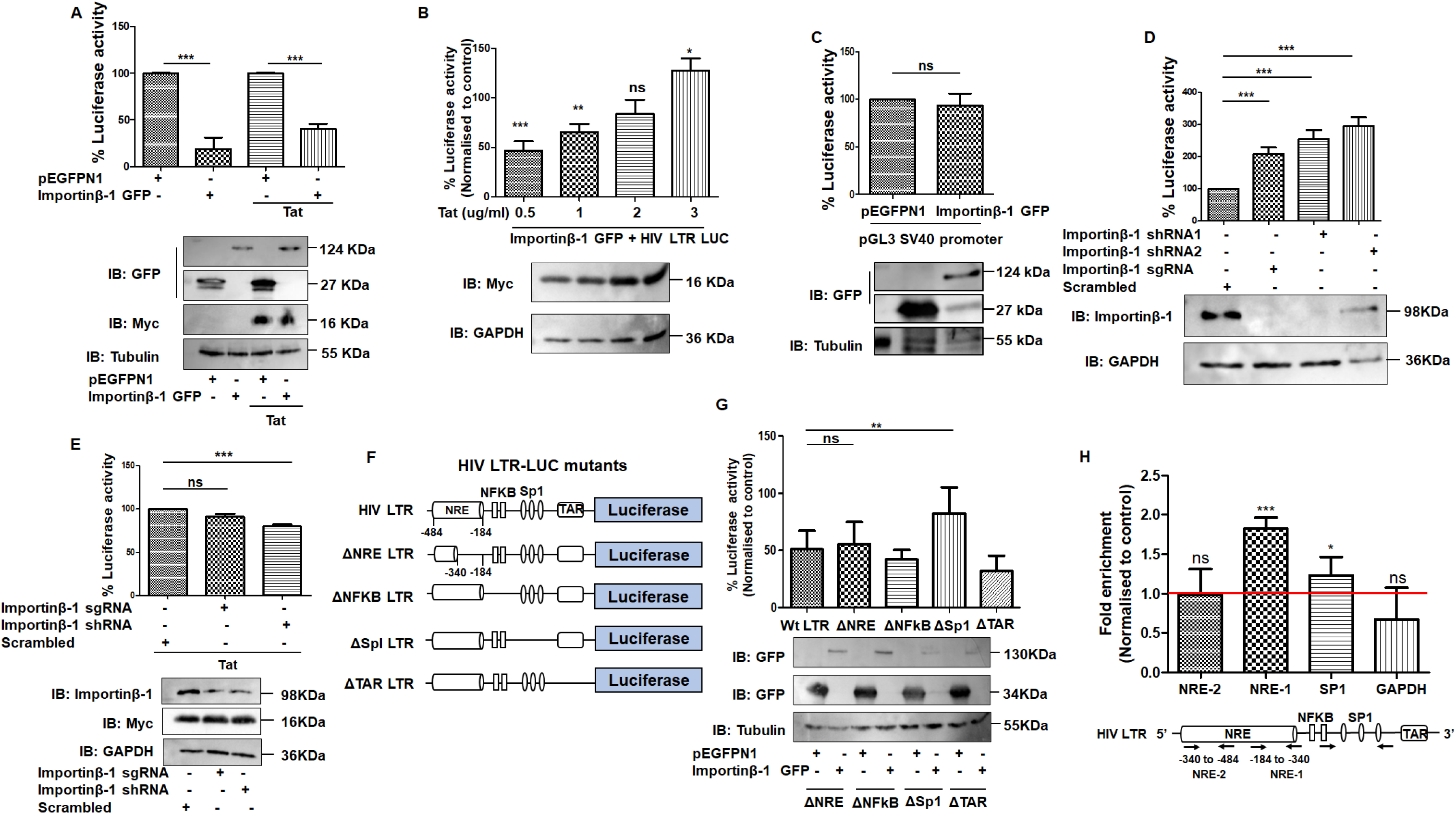
Importinβ-1 specifically inhibits HIV-1 LTR. **(A)** LTR-driven luciferase activity in HEK293T cells upon co-transfection of LTR-LUC and Importinβ-1 GFP or vector control in the presence and absence of pCTat. A corresponding immunoblot is showing the expression of Importinβ-1 GFP and GFP, Tat for these experiments. Tubulin was used as a control. **(B)** LTR-driven luciferase activity in HEK293T cells upon co-transfection of LTR-LUC and Importinβ-1 GFP(1ug/ml) with increased concentration of pCTat. A corresponding immunoblot is showing the expression of Importinβ-1 GFP and GFP, Tat for these experiments. Tubulin was used as a control. **(C)** SV40 viral promoter driven luciferase activity in HEK293T cells upon co-transfection of pGL3-promoter vector with Importinβ-1 GFP or vector control. A representative immunoblot is provided showing the expression of Importinβ-1 GFP and GFP for these experiments. Tat, Tubulin was used as a control. **(D)** LTR-driven luciferase activity in HEK293T cells upon co-transfection of LTR-LUC with Importinβ-1 specific shRNA, sgRNA. A corresponding immunoblot showing the expression of Importinβ-1, GAPDH was used as a control. **(E)** LTR-driven luciferase activity in HEK293T cells upon co-transfection of LTR-LUC along with Importinβ-1 specific shRNA, sgRNA and pCTat. The corresponding immunoblot showing the expression of Importinβ-1, Tat, GAPDH was used as a control. **(F)** Schematic representation of chimeric LTR-LUC mutants. **(G)** LTR-driven luciferase activity in HEK293T cells upon co-transfection of LTR-LUC and its mutants with Importinβ-1 GFP or vector control. The corresponding immunoblot showing the expression of Importinβ-1 GFP, GFP, Tubulin was used as a control. **(H)** qRT-PCR was performed with ChIP samples to check the Importinβ-1 binding on HIV LTR and each Bar graph representing the Importinβ-1 occupancy on various sites of HIV LTR. All experiments were done at least 3 times. The significance is determined using an unpaired student’s t-test. The p values are denoted as *** p <0.001; ** p <0.01; * p <0.05, while non-significant values are denoted by n.s.

To corroborate the negative regulation of Importinβ-1 on HIV-1 LTR, the LTR promoter strength was evaluated in the background of Importinβ-1 knockdown and knockout by shRNA and sgRNA respectively in HEK293T cells. Confirming our previous results, we observed an increase in LTR activity upon knockdown and knockout of Importinβ-1 (Figure 6D), which remained unchanged upon Tat over-expression (Figure 6E). We next investigated which region of LTR is crucial for Importinβ-1 mediated inhibition. For this, we made different deletion mutants of HIV LTR (ΔNRE (Negative Regulatory Region deleted -184 to -340), ΔNF-κB, ΔSP1, ΔTAR) and assayed for luciferase activity (Figure 6F). The reporter assays indicated that Importinβ-1 reduced LTR activity through SP1 site, as reflected by loss of Importinβ-1 inhibitory activity upon deletion of this region (Figure 6G). Next, we have generated individual SP1 sites deletion in LTR and analysed the Importinβ-1 mediated inhibition (Figure S8C). We have observed that even single SP1 site is sufficient for Importinβ-1 mediated inhibition (Figure S8D). We also performed ChIP assays using anti-GFP antibody. The ChIP assay showed enrichment of NRE-1 (-184 to -340) region suggesting possible direct or indirect interaction of Importinβ-1 to this region of LTR. It was interesting to note that SP1 site showed marginal enrichment. Overall, we infer that Importinβ-1 inhibited HIV replication by negatively regulating HIV-1 LTR activity via SP1 and NRE sites (Figure 6H).

We next searched for the domain of Importinβ-1 responsible for regulating LTR activity. Towards this truncated mutants of Importinβ-1 were generated. We deleted RanGTP, IAB binding domain (named ΔRanGTP), N-terminal region from 1-280 amino acids (ΔN-Term), and C-terminal 500 to 860 amino acids (ΔC-Term) regions (Figure 7A). The cellular localizations of these deletion mutants were checked microscopically, and it was observed that all mutants were present both in the nucleus and cytoplasm in presence of pNL4.3 (Figure 7B). We observed that with the deletion of either N-terminal or C-terminal regions, Importinβ-1 lost its inhibitory activity, increasing intracellular p55 and released p24 (Figure 7C, D). Next, we checked the effect of these Importinβ-1 mutants on HIV full-length LTR reporter construct in the presence and absences of Tat. Similar to what was observed upon pNL4.3 transfection, ΔN-Term, and ΔC-Term lost their inhibitory impact on HIV LTR (Figure 7E, F), corroborating the inference that Importinβ-1 inhibited HIV LTR through its N and C-terminal domains.

**Figure 7:**
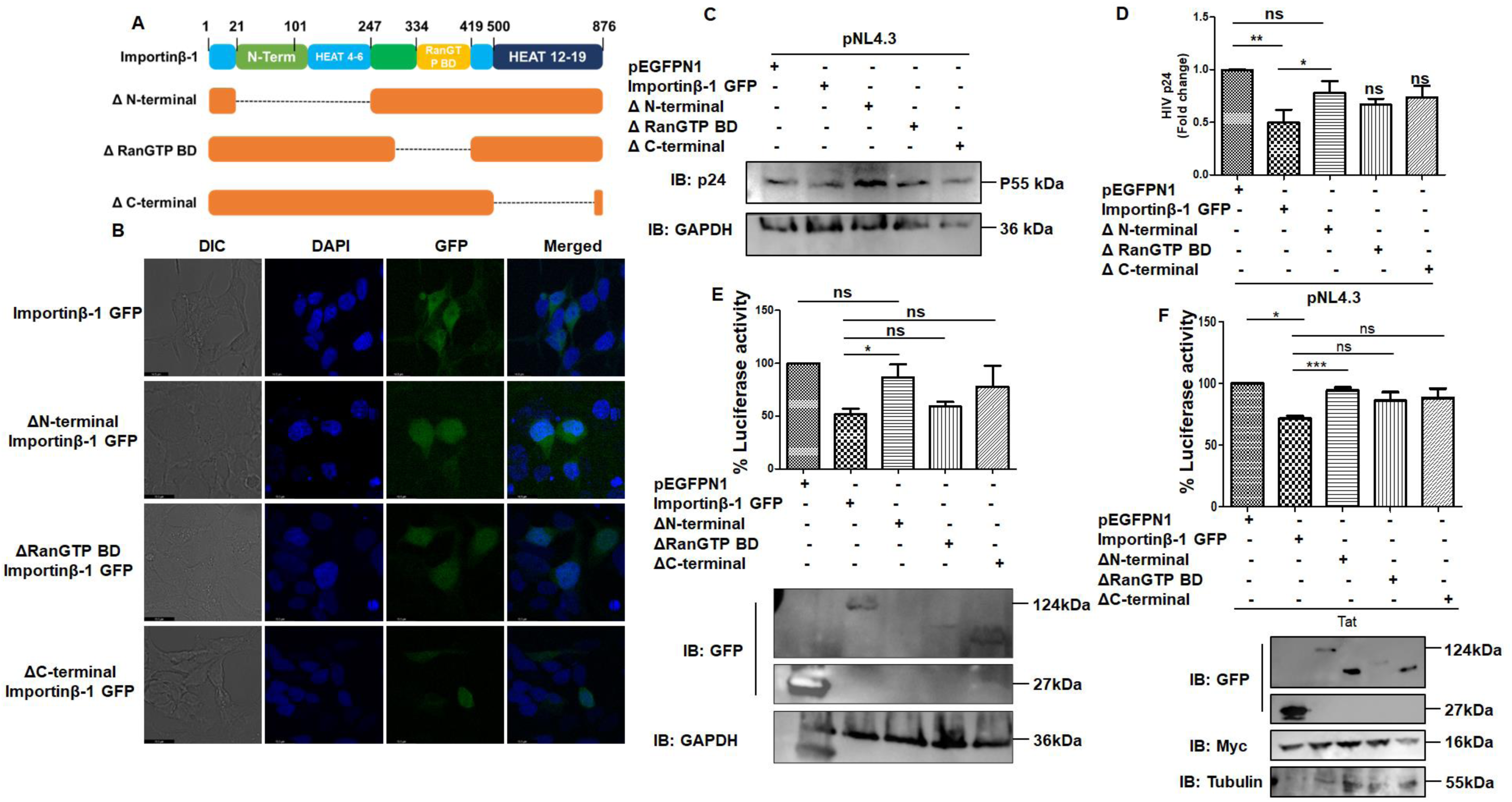
N-terminal and C-terminal regions of Importinβ-1 is important for regulating at HIV LTR. **(A)** Schematic representation of truncated mutants of Importinβ-1 domains. **(B)** Confocal microscopy for subcellular localization of Importinβ-1 GFP and its truncated mutants in HEK293T cells upon co-transfection with pNL4.3. **(C)** Representative immunoblot showing the intracellular p55 levels in HEK293T cells upon co-transfection of pNL4.3 with Importinβ-1 GFP or its truncated mutants. **(D)** Levels of released virus determined by p24 ELISA in supernatant of HEK293T cells upon co-transfection of pNL4.3 with Importinβ-1 GFP or its truncated mutants. **(E)** LTR-driven luciferase activity in HEK293T cells upon co-transfection of LTR-LUC with Importinβ-1 GFP or its truncated mutants. The corresponding immunoblot showing the expression of Importinβ-1 GFP and GFP, GAPDH was used as a control. **(F)** LTR-driven luciferase activity in HEK293T cells upon co-transfection of LTR-LUC with Importinβ-1 GFP or its truncated mutants, in presence of Tat. The corresponding immunoblot showing the expression of Importinβ-1 GFP, Tat and GFP, Tubulin was used as a control. All experiments were done at least 3 times. The significance is determined using an unpaired student’s t-test. The p values are denoted as *** p <0.001; ** p <0.01; * p <0.05, while non-significant values are denoted by n.s.

Combining all the observations, we propose that Importinβ-1 promotes viral core entry during pre-integration stages where basal levels of endogenous Importinβ-1 and packaged Importinβ-1 in virions are supportive. However, it inhibits HIV replication by acting as a potential restriction factor by regulating HIV-1 LTR activity through NRE and SP1 regions during post-integration stages of viral life cycle (Figure 8).

**Figure 8:**
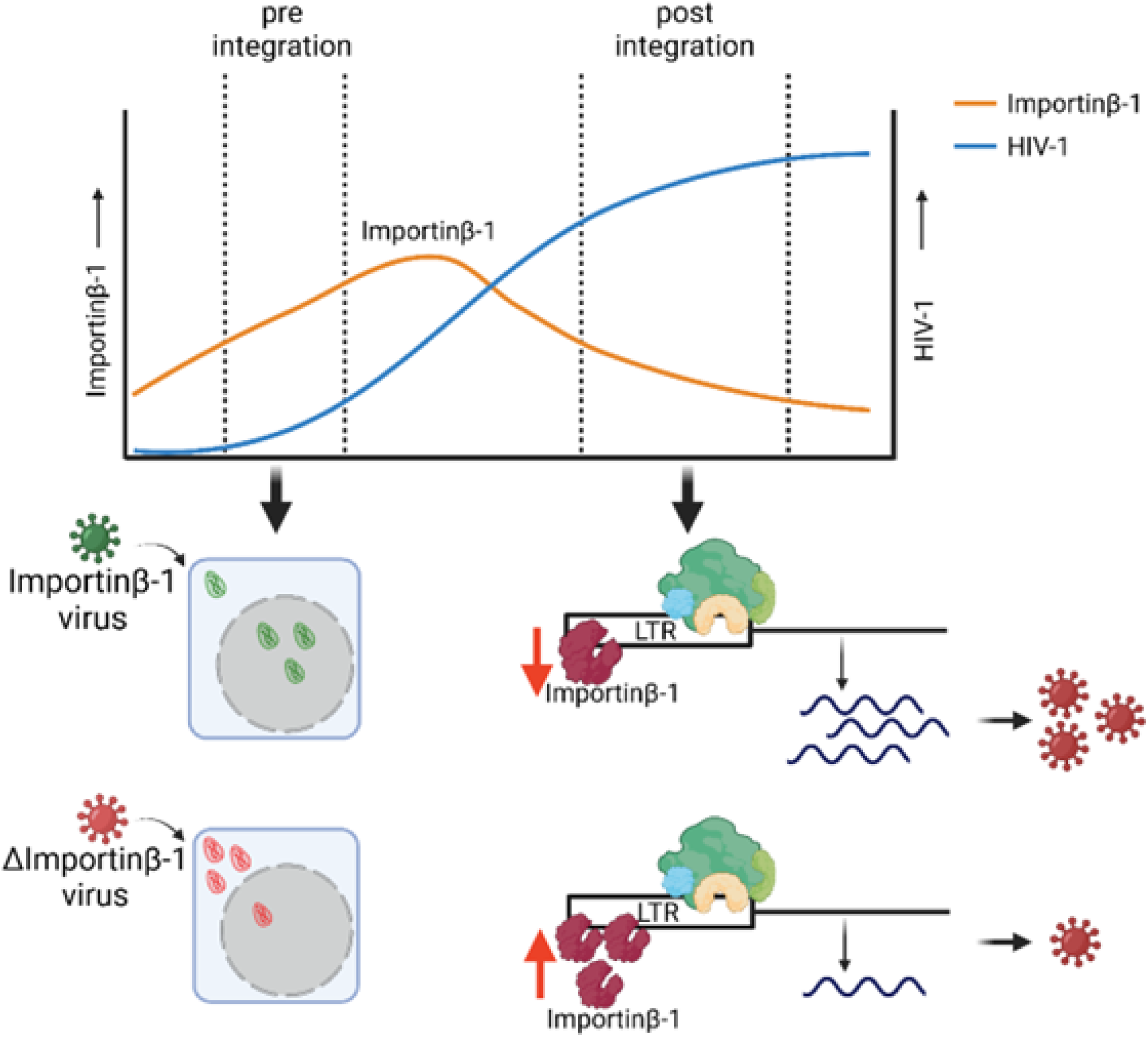
Proposed illustration of Importinβ-1 mediated stage specific regulation on HIV replication. The host Importinβ-1 protein is encapsidated into viruses egressing from immune cells but not the viruses from astrocytes. The packaged Importinβ-1 assists the nuclear entry of the viral core even under low levels of endogenous Importinβ-1 during pre-integration stage. On the contrary, high levels of endogenous Importinβ-1 reduce LTR-driven viral transcription, thereby reducing viral propagation. Sequestering of endogenous Importinβ-1 into emerging virions is an escape strategy adopted by HIV from the inhibitory impact of Importinβ-1. Such stage-specific regulation imposed by the host factors and consequent sequestration of such factors from the cells supports the concept of evolutionary conflict and adaptation between viruses and their hosts.

## Discussion

HIV population has immense diversity with varied pathogenesis, transmission, immune escape, and clinical management [24]. Evolution of viral genome and broad host cells tropism are the critical factors for the diversity and generation of HIV quasispecies within a host. In our study we noticed that virus which emerges from the HEK293T, CD4+T lymphocytes have more infectivity than the virus which emerges from astrocytes. This is to be noted that there is not much genetic diversity in these viruses that we used, but differences in the infectivity were very prominent. Mass spectrometry analysis of virions from different producer cells revealed several host factors that are differentially packaged. With the viruses from astrocytes compromised in infectivity, when compared to those from CD4+ T lymphocytes, in this study we focused on understanding in detail the impact of Importinβ-1, which was encapsidating in the viruses which emerged from HEK293T, and CD4+T lymphocytes but not from the astrocyte cells. Earlier, Santos S, et al also detected Importinβ-1 in the mass spectrometry analysis in HIV core derived from the non-activated THP-1 cells [25]. Interestingly, we did not observe encapsidation of GFP-tagged Importinβ-1 upon over-expression, although we noticed increased infectivity of the viruses egressing from GFP-tagged Importinβ-1 over-expressing 1321N1 and HEK293T cells (data not shown).

Nuclear entry is the essential step for HIV infection and previously it has been reported that many of the host proteins like Nucleoporins (NUPs), NUP358, NUP153 interacted with the incoming HIV-1 Capsid, facilitating the nuclear import of the viral core [26,27]. Similarly, Cleavage and Polyadenylation Specificity Factor 6 (CPSF6), soluble nuclear transport receptors (NTRs) of Transportin-1, Transportin-3 are involved in the nuclear import of the HIV core [28,29]. These factors although have been shown to interact with the viral core/Capsid and induce uncoating, these have not been shown to be directly involved in import of HIV core. Recent studies showed that KPNA2 confers the nuclear import of Pre-Integration Complex (PIC) by interacting with the Capsid protein [6]. It is to be noted that KPNA2 itself depends on the Importinβ-1 for the nuclear import. Our study clearly showed that Importinβ-1 encapsidated in the budding virus increased import of the viral core to the nucleus. Further we have shown that Importinβ-1 interacted with Gag and Capsid proteins for its encapsidation. Also, earlier findings indicated that Capsid binding FG motifs are conserved in many unrelated proteins present in the cytoplasm [30]. Importinβ-1 has two FG repeats at C-terminal, which strengthen the reason of encapsidation of Importinβ-1 in the virion by interacting with the Capsid. Additionally, our data also provided evidence that encapsidated Importinβ-1 is important for the nuclear import in the Importinβ-1 deficient cells. These observations suggested that Importinβ-1 plays a critical role in nuclear-entry of viral core.

Another property of Importinβ-1 that we noticed is its ability to regulate gene expression. We observed that it regulates HIV-1 LTR activity. The negative impact of Importinβ-1 was mediated via NRE and SP1 sites on HIV-1 LTR. Interestingly, Importinβ-1 is also the factor that helps in Importing the positive factors of HIV transcription, such as viral factor (Tat) and host factors (NFκB, and SP1) [31][32][23]. Further, we observed that the inhibitory impact of Importinβ-1 on was restricted to HIV LTR and no such inhibitory impact was observed with SIV or SV40 promoter. Interestingly, our data has shown that ΔRanGTP mutant did not have any inhibitory impact on LTR activity. It is to be noted that import activity of Importinβ-1 is dependent on hydrolysis of RanGTP [33]. A deletion of this mutant makes the regulatory property of Importinβ-1 independent of its import property. This is intriguing as Importinβ-1 is involved in Importing both positive and negative factors for HIV. Hence, we conclude that the regulatory impact on HIV-1 LTR is not because of its import activity. Interestingly, with our ChIP-PCR data showing enrichment of NRE and SP1 regions, one may also speculate that interaction of Importinβ-1 with DNA is either direct or indirect. Although Importinβ-1 has not been shown to interact with DNA, when we performed *in-silico* analysis, it revealed that it has a possible DNA binding pocket and it has higher probability of binding to SP1 region in the LTR (Figure S9, Table S3). The Importinβ-1 relation with DNA binding requires independent and in-depth characterization to understand if specific nucleic acid motifs are preferred by Importinβ-1 to bind.

A major observation that opens several new avenues comes from Importinβ-1 knockdown and knockout experiments which pointed out that while both the complete absence or high expression of Importinβ-1 are detrimental to HIV life cycle, low levels of Importinβ-1 is highly beneficial for HIV. This may be because upon excess or complete depletion of Importinβ-1, the system is compromised in Importing positive factors, such as, Tat, NFκB, and SP1 proteins or unable to help viral core entry into the nucleus. While excess of Importinβ-1 tends to associate with HIV LTR directly or indirectly, thereby limiting viral transcription. With low levels of Importinβ-1 in astrocytes, the virus utilizes the same to enter the nucleus, but in CD4+ T lymphocytes, the high levels of intracellular Importinβ-1is removed by packaging into new virions. The next pertinent question is to understand the mechanism of Importinβ-1 mediated transport of viral core into the nucleus. In addition, we observed that Importinβ-1 levels were elevated upon treatment of cells with cytokines such as IL4, and IFNγ. This prompts further investigation of the role of Importinβ-1 as a cytokine-induced gene and should be studied for its role in innate immune responses against a battery of infections. We envisage a more general phenomenon where Importinβ-1 plays a role in innate immune mechanisms against viruses by modifying gene expression through its DNA associations and regulating nuclear import of host and viral factors.

## Methods

The experiments were performed as per the approved protocols by the institutional Biosafety committees of the University of Hyderabad, Hyderabad.

### Cell lines and antibodies

DMEM (HIMEDIA Cat: AL066A-500ML) was used for maintaining HEK293T, 1321N1, and TZM-bl cell lines, and RPMI-1640 (HIMEDIA Cat: AL162A-500ML) for SUP-T1 cell line. The cell lines were maintained at 37^0^C with 5% CO2 [34], supplemented with 10% Fetal Bovine Serum (Gibco USA cat#:10270-106) and penicillin (100 U/ml/streptomycin (100ug/ml) (HIMEDIA cat#: A001A).

Primary antibodies anti-p24 antibody (Ab9071, Ab63598), Anti-GAPDH (Sc47724), anti-Integrase (sc-69721) and anti-GFP antibody (Sc-9996) were procured from Santa Cruz Biotechnology; anti-KPNB1 (Importinβ) antibody (Ab2811) and anti-Tubulin antibody from ABclonal (AC008) or Sigma (T8328-25UL).

Secondary antibodies were procured from Cell Signalling [anti-mouse IgG HRP, (08/2017)], Santa Cruz Biotechnology [anti-Rabbit IgG HRP, (Sc-2357)] and Abcam [Anti-mouse IgG Alexa 647 (Ab150115)]

### Isolation of primary CD4+T lymphocyte

PBMCs were isolated from four individual healthy donors by using the Histopaque (10771-500ML) density gradient method as described earlier[35]. Around 5 x 10^7^ cells/mL PBMCs were used for isolation of CD4+T lymphocytes by Human CD4+ T Cell Isolation Kit (EasySep™ Cat:17952). 50 ul/mL of the cocktail was added and incubated for 5 min at room temperature. Subsequently, 50 ul/mL of RapidSpheres beads were added to the tubes, placed in the magnetic separator, and incubated for 3 min. Further, the tube together with the magnetic stand was inverted to decant the unbound cells in the supernatant in a fresh tube. The supernatant with CD4+T lymphocytes, thereby, was collected in the fresh tube for further experiments.

### Plasmids

The plasmid pEGFP-N1-Importinβ (Plasmid #106941) was procured from Addgene. This plasmid has been henceforth referred to as Importinβ-1 GFP. The mutant constructs ΔN-term-Importinβ-1, ΔC-term-Importinβ-1, ΔRanGTP-Importinβ-1, ΔNRE-LTR luciferase, ΔNFKB-LTR luciferase, ΔSP1-LTR luciferase, ΔTAR-LTR luciferase, constructs were generated by site-directed mutagenesis using the respective primers listed in (Table S2) with pEGFP-N1-Importinβ or wt LTR-LUC reporter as template respectively. All the constructs were confirmed by sequencing. HIV-1 Rev-Myc was cloned in pLVX-IRES-Puro [36]. The sgRNA targeting Importinβ-1 is designed using the GPP web portal and cloned into the lentiCRISPR v2 vector. The pLKO shRNA targeting Importinβ-1 and scrambled shRNA were obtained from the ShRNA Resource Centre (Indian Institute of Science, Bangalore, India).

### Transfection and infection

HEK cells were seeded 12 h before transfection. Fresh media was added to the cells before transfection. Lipofectamine 2000 (ThermoFisher Scientific, USA), and calcium phosphate precipitation methods were used to transfect HEK293T, 1321N1 cells as previously described [37].

HIV-1 virus particles (NL4.3) were produced by transient transfection of HEK293T cells with pNL4.3 using the calcium phosphate method. The culture supernatant was collected after 48 h of transfection, filtered through a 0.45 µM syringe filter (PALL Cat: 4614), and precipitated by polyethylene glycol (PEG) (Sigma Cat: p6667-1kg, 81260-1kg). For infection of SUP-T1/Primary CD4+T lymphocytes, the cells were pre-treated with 10 ug Dextran (Sigma Cat: D9885-10G) followed by infection with NL4.3 virus. 50 ng/ml or 100 ng/ml of p24 equivalent virus was used to infect cells. After 6 h of infection, cells were thoroughly washed with phosphate-buffered saline (PBS) to remove free virion particles, and cell culture media was replaced with fresh media as described earlier [34]. Viruses were collected from HEK293T, SUP-T1, and 1321N1 48 h post-infection and quantified using a p24 ELISA kit (Advanced Bioscience Laboratory Inc, USA. Cat: 5447). Further, these viruses were used to infect TZM-bl cells at different concentrations, and beta-galactosidase assays were evaluated at various time points. The Beta-galactosidase assay was performed.

### Proteomics Methodology

Tryptic digested and desalted peptides using (C18 ZipTips (Millipore) samples were acquired using Orbitrap Fusion mass-spectrometer (Thermo Scientific™) coupled with EASY-nLC 1200 (Thermo Scientific™). All the acquired data were analyzed using Thermo Proteome Discoverer software (version 2.2.0). Peptides were separated using a 140 min gradient of 5 to 95% phase B (80% acetonitrile and 0.1% formic acid) at a flow rate of 300 nL/min, using EASY-Spray nano flow column (50 cm×75 μm ID, PepMap C18). The parameters used in the MS-MS method are as follows, Orbitrap analyser has a resolution of 60,000, with automatic gain control (AGC) target of 4×10^5^. The scan sequence began with MS1 spectrum from mass range 375–1700 m/z and maximum injection time of 50 ms. MS2 precursors were fragmented by high energy collision-induced dissociation (HCD) and analyzed using the Orbitrap analyser (NCE 35; AGC 5×10^4^; maximum injection time 22 ms, resolution 15,000 at 200 m/z). The proteome identification was performed through the Sequest HT database with 1% FDR and 1 missed cleavage permissible limits as input parameters. Database was searched against the Homo Sapiens and HIV Uniprot database. Total protein level precursor ion tolerance was set at 10 ppm. The precursor and product ion tolerance were set as 10 ppm and 0.05 Da respectively. Carbamidomethylation of cysteine residues (+57.021 Da) was set as static modifications, while oxidation of methionine residues (+15.995 Da) and acetylation of protein N-terminus (+42.011 Da) were set as a variable modification. Peptide-spectra matches (PSMs) were adjusted to a false discovery rate (FDR) value of 0.01. PSMs were identified, and narrowed down to a 1% peptide FDR and then further to a final protein-level FDR of 1%. The protein list ratios were exported for further analysis to Microsoft Excel.

### Generation of Importinβ-1 knockdown and knockout cell lines

The protocol for the generation of knockout and knockdown was followed as described earlier [36]. Briefly, HEK293T cells were seeded in a 6-well plate 12 h before transfection. Transfection was carried with a cocktail of helper plasmid (pMD2.G, 250ng), envelope plasmids (psPAX2,1000ng), lenti-viral vectors (shRNA scrambled, shRNA Importinβ-1, sgRNA Importinβ-1) using calcium phosphate precipitation methods. The lentivirus was collected by filtering the supernatant through a 0.45μm syringe filter after 48 h of transfection. The engineered lentivirus was transduced to the target cells and changed the medium after 12 h of transduction. After a further incubation of 48 h, the cells were selected by increasing Puromycin concentration beginning with 1µg/ml and increasing to 2µg/ml and then 3µg/ml at an interval of 24 h. The Puromycin selected cells were grown in an antibiotic-free medium for 24 h and the Knockdown or Knockout of the gene was assessed by immunoblotting.

### Separation of nuclear and cytoplasmic fractions

3 x 10^6^ TZM-bl cells were washed thrice with 1XPBS and resuspended in cytoplasmic extraction (CE) buffer (10 mM HEPES pH 7.6, 60 mM KCl, 1 mM EDTA, 0.075 % NP-40, 1 mM DTT, 1 mM PMSF). The samples were gently pipetted two or three times and incubated on ice for 6 min, followed by centrifugation at 1000 rpm for 5 min at 4^0^C. The supernatant containing cytoplasmic fraction was collected into a fresh tube. The pellet containing the nuclear fraction was washed twice with CE buffer excluding NP-40. Both the cytoplasmic and nuclear fractions were processed for western blot analysis.

### Western blotting

The cell lysates were prepared using RIPA buffer containing PIC. The cell extracts were fractionated on SDS-PAGE and transferred onto nitrocellulose membrane (PALL, BioTrace^TM^ NT nitrocellulose, Cat: 66485) by semi-dry electrophoretic transfer at 15V for 20 min. Protein transfer on the membrane was visualized by staining with Ponceau solution (Fluka Analytical, Cat: 81460-25g). After blocking with 5% of skimmed milk (HIMEDIA, Cat: GRM1254-500G) for 1 h at room temperature, primary antibody (as per recommended dilution) was added and incubated overnight at 4°C. The following day, the blot was washed with 1X PBST, and the appropriate secondary antibody (as per recommended dilution) was added for 2 h at room temperature (RT). The blot was developed using a WesternBright^TM^ ECL kit (Advansta, K-12045-D20) and visualized using a Bio-Rad ChemiDoc instrument.

### Luciferase assay

HEK293T were transfected with the reporter plasmids and maintained for 48 h and in the case of TZM-bl cells, they were infected with appropriate viruses and maintained different time points respective to the type of viruses used. The cells were washed with cold 1X PBS (Cat: HIMEDIA, TS1099-20L) and collected after trypsin treatment. Cells were centrifuged at 1000 rpm for 5 min and washed with PBS. Cells were lysed in Reporter lysis buffer (Promega, Cat: E397A). The samples stored at -80°C were thawed and kept at RsT for 20 min. Freeze-thaw was repeated 2 times. The samples were centrifuged at 11000 rpm for 5 min at 4°C and the supernatant was collected in 1.5 ml centrifuge tubes. Firefly Luciferase activity was measured using the Modulus single-tube multimode reader as per the protocol [38,39].

### β-galactosidase assay

TZM-bl cells were infected and maintained until different time points respective to the type of viruses used. The cells were washed with cold 1X PBS (Cat: HIMEDIA, TS1099-20L) and 100µl of lysis buffer (Tris PH 8, 0.5% Triton X100) was added and incubated at 37^0^C for 30 min. After incubation, the sample was pipetted 5 to 10 times and harvested at 3000 rpm for 10 min. 50µl of lysate was aliquoted into a fresh 96 well plate and incubated with 100 ul of substrate solution ONPG (1.5 mg/ml ONPG, 10mM sodium phosphate buffer pH 7.5, 30mM MgCl2, 0.075% v/v β-ME). After adding the substrate solution, the sample was incubated for 30 min at 37 ^0^C, and the reaction was stopped by the stop solution (2N NaCO3). The absorbance was measured at 450 nm in a microplate reader (PowerWave XS2).

### Immunofluorescence microscopy

HEK293T cells were seeded in a 6-well plate one day before transfection with different constructs. After transfection, the cells were washed twice with cold 1X PBS and fixed with 4% formaldehyde (Sigma, Cat: F1635-500ML) for 20 min and subsequently lysed with 0.5% Triton X100 (Qualigens, Cat:10655) for 15 min at RT. The cells were washed twice with PBS and blocked with 3% BSA (SRL, Cat: 83803) for 30 min. The cells were then stained for 2 h in primary antibody (prepared in 1% BSA) at RT. After 3 washes with PBS, cells were incubated for 1 h with the secondary antibody tagged with Alexa 647 (Ab150115) or Alexa 488 (Ab150077). Cells were then washed with PBS, and the nuclei were stained with DAPI (Abcam, Cat: ab104139). Images were captured using a Leica Confocal Laser Scanning microscope.

### Image Quantification

The quantification of microscopic images was done using ImageJ software. The image was uploaded on the ImageJ software and the total intensity of the cell was quantified. The fluorescence intensity of Capsid in the nucleus and cytosol was calculated separately using the ROI tool for DAPI, and Capsid (in nucleus and cytosol). The fluorescence intensity was converted into percentage sub-cellular localization of Capsid using the total intensity of Capsid. The graphs were plotted with the normalized values.

### Co-immunoprecipitation (Co-IP)

HEK293T cells were transfected with pNL4.3 and incubated for 48 h. The cells were lysed with Nonidet P-40 (NP40) buffer supplemented with PIC (Cat: 786-437, G-Bioscience). Importin-ß was precipitated by protein A/G beads conjugated with anti-Importinβ-1 or anti-p24 antibody while incubating for 12 h in 4°C, followed by washes with 1X TBST buffer. The bound protein was eluted by boiling in 2X loading buffer (without reducing agent) at 100°C for 10 min. The beads were separated and proceeded with immunoblotting to identify interacting partners.

### Two LTR circles measurement

To measure the nuclear entry of the virus, two LTR circles were measured. Total DNA was isolated from the infected TZM-bl cells using the GeneJET Genomic DNA purification kit (Thermo Scientific, Cat: K0721) as per the manufacturer’s protocol. An equal concentration of the isolated DNA was used for qRT-PCR analysis by using the primers of 5’LTR and 3’LTR (Table S2).

### RNA extraction and qRT-PCR

To measure the mRNA levels, total RNA was isolated from the cells using a GeneJET RNA purification kit (Thermo Scientific, Cat: K0731) according to the manufacturer’s protocol. 10 µg of isolated RNA was treated with DNase I (NEB, Cat: M0303S) to remove any contaminating genomic DNA. 1µg of DNase I treated RNA was converted into cDNA using an iScript cDNA synthesis kit (BIO-RAD, Cat:1708891) according to the manufacturer’s protocol. The cDNA was diluted 10 times with nuclease-free water as the qRT-PCR template. qRT-PCR was performed using the iTaq Universal SYBR green Supermix (BIO-RAD, Cat:1725121). One cycle of denaturation (95°C for 30 sec) was followed by 40 cycles of amplification (95°C for 15 sec, 55°C for 30 sec, and 60°C for 20 sec). The total reaction was prepared in a 96-well plate (applied biosystems, Cat: 4346907) compatible with the Applied Biosystems qRT-PCR instrument. The applied biosystems Quantstudio ^TM^ 3 real-time PCR system was used for the qPCR studies.

### DNA-Protein Docking

HDOCK server was used for the prediction of protein-DNA docking. We used Importinβ-1 as a receptor and dsDNA corresponding to NFκB and SP1 binding sites on HIV-1 LTR, were used as ligands. The structure of Importinβ-1 (PDB ID: 1QGR) was obtained from the Protein Data Bank (https://www.rcsb.org/). The dsDNA structures were prepared using build and edit tool in discovery studio 3.5 for each given sequence and DNA model was docked to the free conformations of Importinβ-1 protein. Both protein and DNA were defined to be semi-flexible on all their length after the rigid-body docking stage. The DNA-protein docking was performed using blind docking protocol where the entire protein was selected in grid mapping covering all possible binding sites.

### DNA Immunoprecipitation (ChIP)

To check the interaction between Importinβ-1 and DNA, HEK293T cells were co-transfected with LTR luc and Importinβ-1 GFP construct. Cells were harvested after 48 h post-transfection and crosslinked with 0.75% of formaldehyde (Sigma, Cat: F1635-500ML) for 10 min at room temperature with mild shaking. This was followed by the addition of 125 mM Glycine (SRL, Cat: 66327) for 5 min at RT with mild shaking. The cells were washed twice with PBS, and then sonication for 5 min at 4°C to shear the DNA. The sonicated samples were centrifuged (Eppendorf centrifuge 5417R) at 11000 rpm (11228 rcf) for 5 min at 4°C. The supernatant was collected and 2 µg of anti-GFP antibody pre-conjugated with protein AG beads (sc2003) was added and immunoprecipitated overnight. Subsequently, the beads were washed five times with RNA immunoprecipitation buffer (Abcam RNA immunoprecipitation protocol), followed by RNase and protease treatment, DNA isolation, and qRT-PCR with specific primers (Table S2).

### Statistical Analysis

All experiments were performed at least three times. The error bar representing the standard error mean and statistical significance was determined by using the P value, the P value <0.05 was considered statistically significant.

## Supporting information

Mass spectrometry analyses

## Acknowledgments

The study is funded by SERB POWER (SPG/2021/002627) and DBT (BT/HRD/NWBA/38/09/2018) to SB. We also acknowledge SERB SUPRA support (SPR/2021/000137) to SB, UGC JRF-SRF to SY, and DBT JRF, PMRF to SS, and DST-WISE-PDF support to SSx (DST/WISE-PDF/CS-22/2023). We thank Dr. Reddy’s Institute of Life Sciences, Hyderabad for gifting HEK293T. We thank Prof. Anand Kumar Kondapi, UoH, for sharing 1321N1 cells, kind gift of pNL4.3 proviral DNA clone and permitting us to use their cell culture facility. Prof. Uday Kumar Ranga, Jawaharlal Nehru Centre for Advanced Scientific Research (JNCASR) for TZM-bl cells and Dr. S. Jameel, ICGEB, New Delhi for SUP-T1. We thank Dr. Radha Chauhan for sharing the anti-Importinβ-1 antibody. Wt LTR, ΔSP1 -III, ΔSP1-III+II were gifted by Dr. Frank Kirchhoff from Ulm University Medical Center, Ulm, Germany [40]. We also thank Ms. Deepti for helping with confocal microscopy imaging, DST-FIST to the Department of Biochemistry, the DBT-BUILDER grant to the School of Life Sciences (BT/INF/22/SP41176/2020), and the Institution of Eminence supported projects RC1-20-017 and RC4-21-012 to SB and IoE to the University of Hyderabad MHRD (F11/9/2019-U3(A) for infrastructure support to the department of Biochemistry, School of Life Sciences and University common facilities are acknowledged. We also thank UoH-NIAB BSL3/ABSL3 facility. We recognize GraphPad prism for generating graphs and statistical analysis.

## Data Availability Statement

This study includes no data deposited in external repositories. All relevant data can be found in the manuscript and supporting information files.

## Author Contributions

SY: Conceptualization, investigation, designed and performed experiments, analysed data, manuscript writing (original draft), and editing. SS: investigation, performed experiments, analysed data. SSx: Performed *Insilco* experiments and manuscript writing. VN, SR: mass-spectrometry, peptide recognition and analysis. SB: Conceptualization, supervision, review and editing manuscript, funding acquisition. All the authors read the final draft and approved.

## Supporting information

The following are available online: Supplementary File 1 (Table S1, Table S2 and Figure S1-S9, Table S3), Supplementary File 2 (Analysed Mass spectrometry data sheets).

