## Supplementary Figure file 1 for "The stage-specific regulation imposed by Importinβ-1 on HIV-1 propagation and infectivity dynamics"

**Table S1. Comparative mass spectrometry data analysis in the HEK293T_NL4.3, SupT1_NL4.3 and 1321N1_NL4.3 virus.**

| Name of the protein | HEK293T | SUP-T1 | 1321N1 | Function |
|---|---|---|---|---|
| Importin Beta-1 | + | + | - | Import of proteins to nucleus |
| Elongation factor 1-gamma | - | + | + | Responsible for the enzymatic delivery of aminoacyl tRNAs to the ribosome |
| Seryl -tRNA synthase | + | + | - | Involved in protein translation |
| RNA helicase (DDX3X) | + | - | - | Binds RNA G-quadruplex (rG4s) structures, including those located in the 5'-UTR of NRAS mRNA |
| Serine/threonine-protein phosphatase 2A catalytic subunit beta isoform | + | + | - | PP2A can modulate the activity of phosphorylase B kinase casein kinase 2 |
| T- complex protein-1 | + | - | + | assists the folding of proteins upon ATP hydrolysis |
| HIV-Polyprotein | + | + | + | Viral structural protein |

**Table S2. List of primers used in the study**

| | | |
|---|---|---|
| pCDNA HIV-Tat | FP | 5'ACGAATTCACCATGGAGCCAGTAGATCCTAGACTAG3' |
| | RP | 5'TATCTAGATTCCTTCGGGCCTGTCGGGTCC3' |
| 2LTR circles | FP | 5'AACTAGGGAACCCACTGCTTAAG3' |
| | RP | 5'TCCACAGATCAAGGATATCTTGTC3' |
| CD4 qPCR | FP | 5'AACTTTCCCCTGATCATCAAGAATCTT3' |
| | RP | 5'CCCTTGGACTCCTACATTGCACTGA3' |
| CD3 Delta Receptor | FP | 5'ACGAGGAATATATAGGTGTAATGGGACA3' |
| | RP | 5'AGCCCCAGACAGCCTTCCAGT3' |
| qPCR spliced | FP | 5'ACGAAGAGCTCATCAGAACAGTCAGAC3' |
| | RP | 5'CAGAAGTTCCACAATCCTCGTTACAAT3' |
| qPCR RRE | FP | 5'GCAGTGGGAATAGGAGCTTTGTTC 3' |
| | RP | 5'GAGCTGTTGATCCTTTAGGTATCTTTCC 3' |
| GAPDH qPCR | FP | 5'TGTTGCCATCAATGACCCCTT3' |
| | RP | 5'CTCCACGACGTACTCAGCG3' |
| ΔNFKB LTR LUC SDM | FP | 5'TTGTTACAAAGGGAGGCGTGGCCTGGGCGG3' |
| | RP | 5' CGCCTCCCTTTGTAACAAGCTCGATGTCAA3' |
| ΔSPI LTR LUC SDM | FP | 5' ACTTTCCAGGAGCCCTCAGATGCTGCATAT3' |
| | RP | 5' TGAGGGCTCCTGGAAAGTCCCCAGCGGAAA3' |
| ΔTAR LTR LUC SDM | FP | 5' CCTCAGATGCCTGGGAGCTCTCTGGCTAGC3' |
| | RP | 5' GCTCCCAGGCATCTGAGGGCTCGCCACTCC3' |
| ΔNRE LTR LUC SDM | FP | 5' CTTACAAGGACCCTGAGAGAGAAGTGTTAG3' |
| | RP | 5' TCTCAGGGTCCTTGTAAGTCATTGGTCTTA3' |
| NRE Chip | FP | 5' ATCTACCACACACAAGGCTACTTCC3' |
| | RP | 5' CCACTCTAACACTTCTCTCTCAGGGT3' |
| SpI ChiP | FP | 5' TTTGACAGCCGCCTAGCATTTC3' |
| | RP | 5' CATCTGAGGGCTCGCCACTCC3' |

| | | |
|---|---|---|
| Non-NRE ChiP | FP | 5’ AGACCTGGAAATAACATGGAGCAAT3’ |
| | RP | 5’ GAGTGAATTAGCCCTTCCAGTCC3’ |
| ΔN Terminal | FP | 5’ CTGGAAGCGTACATGGAGACATATATGGGT 3’ |
| | RP | 5’ CTCCATGTACGCTTCCAGCTCCAGCCGATC 3’ |
| Δ RAN GTP BD | FP | 5’ CAAGGGATAGACGTTGCTGATGATCAGGAA 3’ |
| | RP | 5’ AGCAACGTCTATCCCTTGTAAAGCCACCTC 3’ |
| Δ C Terminal | FP | 5’ TACTGCTTACTTGCTACATGGGCAACAAAA 3’ |
| | RP | 5’ TGTAGCAAGTAAGCAGTAAGTAGCTGGTTC 3’ |
| KPNB1 sgRNA-1 | FP | 5’CACCGCACACAGTGTCCAGATACGA3’ |
| | RP | 5’AAACTCGTATCTGGACACTGTGTGC3’ |
| KPNB1 sgRNA-2 | FP | FP: 5’CACCGAGCTCCTAGAGACTACAGAC3’ |
| | RP | RP: 5’AAACGTCTGTAGTCTCTAGGAGCTC3’ |
| KPNB1 sgRNA-3 | FP | 5’CACCGTCAAGGCACAATATCAGCAG3’ |
| | RP | 5’AAACCTGCTGATATTGTGCCTTGAC3’ |
| LRT | FP | 5’TGTGTGCCCGTCTGTTGTGT3’ |
| | RP | 5’GAGTCCTGCGTCGAGAGAGC3’ |
| Alu-Gag | Alu FP | 5’ GCCTCCCAAAGTGCT GGGATTACAG 3’ |
| | Gag RP | 5’ GCTCTCGCACCCATCTCTCTC C 3’ |
| LTR | FP | 5’ GCCTCAATAAAGCTTGCCTTG A 3’ |
| | RP | 5’ TCCACACTGACTAAAAGGGTCTGA 3’ |

**Supplementary Figure S1.**

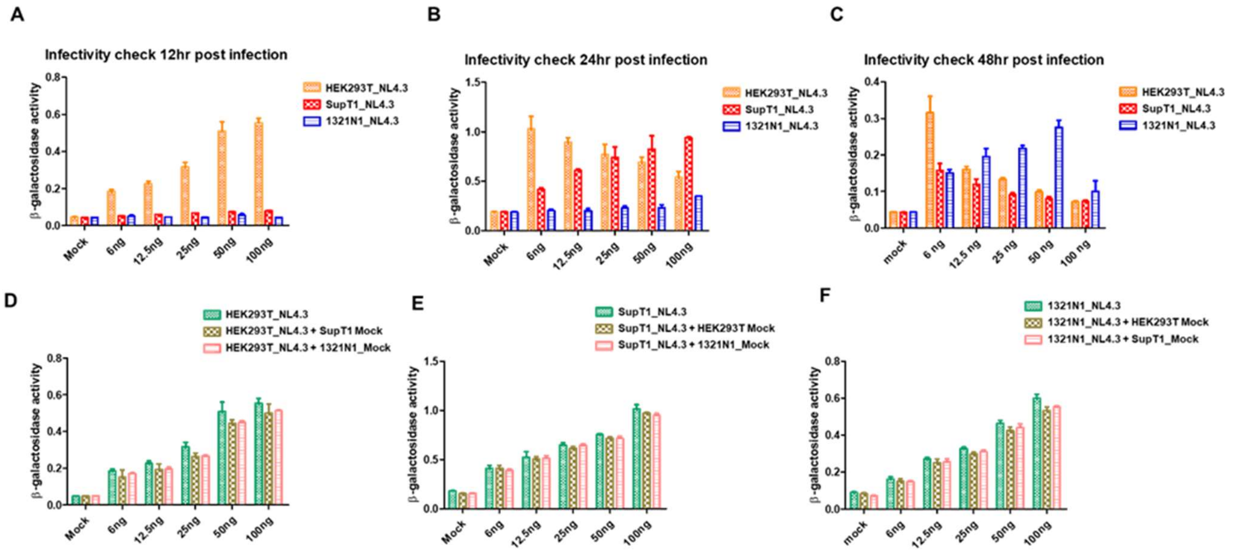


**Figure S1: Differential infectivity of virus, emerges from the different cells.** (**A-C**) LTR driven β-galactosidase activity upon infection of TZM-bl cells with HEK293T_NL4.3, SupT1_NL4.3, 1321N1_NL4.3 viruses (ng/ml) at different time points (HEK293T_NL4.3 12hrs, SupT1_NL4.3 24hrs, 1321N1_NL4.3 48hrs). **(D)** LTR driven β-galactosidase activity upon infection of TZM-bl cells with HEK293T_NL4.3 (ng/ml) in presence of SupT1 mock or 1321N1 mock (100ng/ml) after 12 hours post-infection. **(E)** LTR driven β-galactosidase activity upon infection of TZM-bl cells with SupT1_NL4.3 (ng/ml) in presence of HEK293T mock or 1321N1 mock (100ng/ml) after 24 hours post-infection. **(F)** LTR driven β-galactosidase activity upon infection of TZM-bl cells with 1321N1_NL4.3 (ng/ml) in presence of SupT1 mock or HEK293T mock (100ng/ml) after 48 hours post-infection.

**Supplementary Figure S2.**

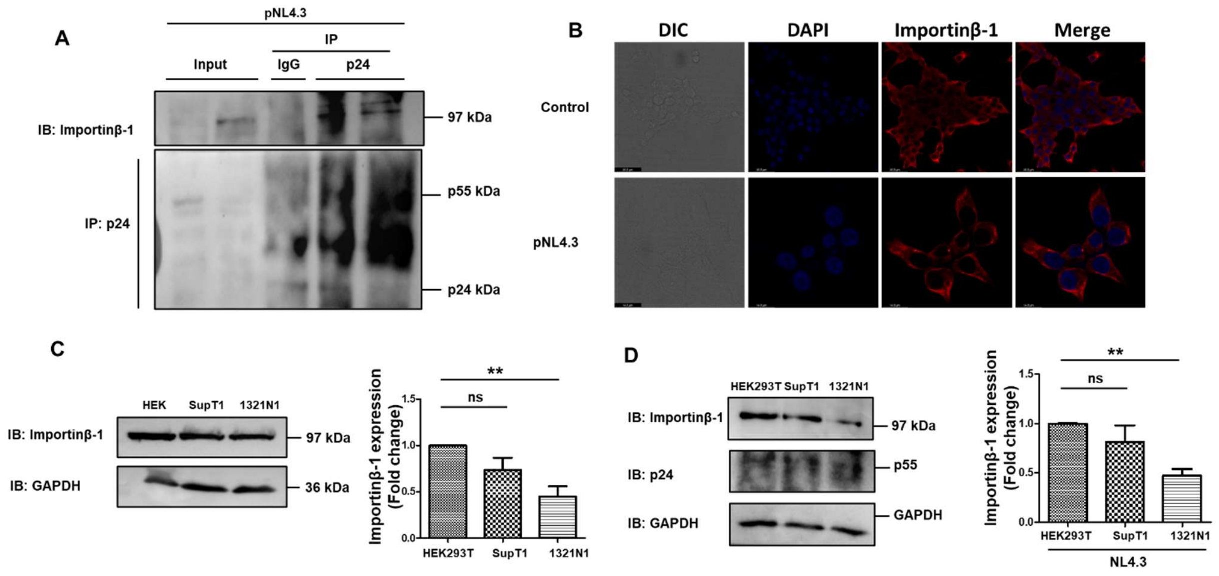


**Figure S2: Interaction, expression, localisation of Importinβ-1 in HIV background. (A)** Co-immune precipitation showing Importinβ-1 interaction with Gag and Capsid proteins upon pNL4.3 transfection in HEK293T cells. Mouse IgG was used as an isotypic control. **(B)** Confocal microscopy for subcellular localisation of endogenous Importinβ-1 in the pNL4.3 transfected and untransfected cells of HEK293T cells. **(C)** Representative immunoblot showing the Importinβ-1 expression in the HEK293T, SUP-T1, 1321N1 cells without infection, besides bar graph representing the quantification of Importinβ-1 expression from three independent of experiment. **(D)** Representative immunoblot showing the Importinβ-1 expression in the HEK293T, SUP-T1, 1321N1 cells after 48hrs of infection, besides bar graph representing the quantification of Importinβ-1 expression from three independent of experiment. All experiments were done at least 3 times. The significance is determined using an unpaired student's t-test. The p values are denoted as ** $p < 0.01$, while non-significant values are denoted by n.s.

**Supplementary Figure S3.**

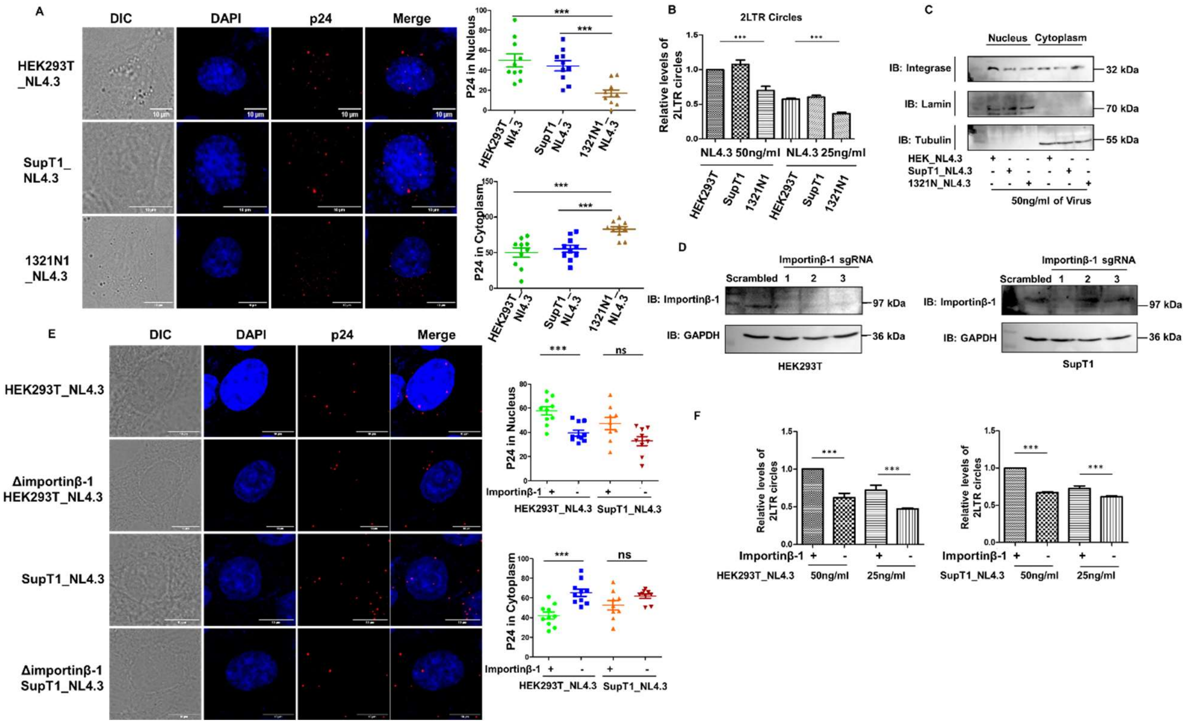


**Figure S3: Importinβ-1 packaging is critical for viral core entry into the nucleus. (A)** Confocal microscopy for sub cellular localisation of capsid protein in the infected TZM-bl cells, 8 hrs of infection. Quantification of capsid protein presence in the nucleus and cytoplasm of infected TZM-bl cells using ImageJ software. **(B)**The bar graph representing the two LTR circles in the infected TZM-bl cells by performing the qRT-PCR with genomic DNA, 12hrs of infection. **(C)** Representative immunoblot showing Integrase in the nuclear and cytoplasmic fractions of the infected TZM-bl cells, after 8 hrs of infection. Lamin and Tubulin were used as a nuclear and cytoplasmic controls respectively. Bar graph showing immunoblot quantification of Integrase in the nuclear and cytoplasmic fractions of three independent experiments. **(D)** Immunoblot showing the Importinβ-1 knockout generation in the HEK293T cells, SUP-T1 cells. **(E)** Confocal microscopy for sub cellular localisation of capsid protein in infected TZM-bl cells with HEK293T_NL4.3, SupT1_NL4.3 and ΔImportinβ-1 HEK293T_NL4.3, ΔImportinβ-1 SupT1_NL4.3 viruses, 8 hrs of infection. Quantification of capsid protein presence in the nucleus and cytoplasm of infected TZM-bl cells using ImageJ software. **(F)** The bar graph representing the two LTR circles in infected TZM-bl cells with HEK293T_NL4.3, ΔImportinβ-1 HEK293T_NL4.3, SupT1_NL4.3, ΔImportinβ-1 SupT1_NL4.3 viruses. The qRT-PCR was performed with genomic DNA, after 12hrs of infection. All experiments were done at least 3 times. The significance is determined using an unpaired student's t-test. The p values are denoted as *** p <0.001, while non-significant values are denoted by n.s.

**Supplementary Figure S4.**

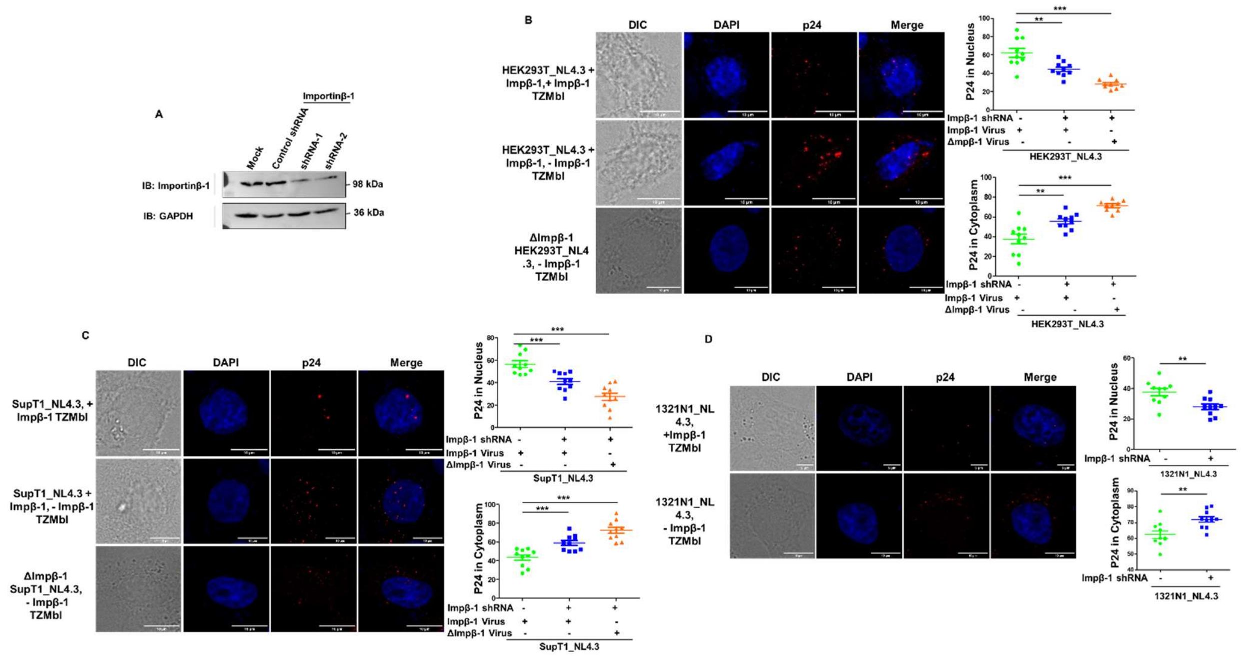


**Figure S4: Packaged Importinβ-1 is sufficient to import the viral core into nucleus in Importinβ-1 depleted cells. (A)** Immunoblot representing the expression of Importinβ-1 in TZM-bl cells after Importinβ-1 specific shRNA expression, GAPDH was used as a loading control. **(B)** Confocal microscopy for sub cellular localisation of capsid protein in the HEK293T_NL4.3 +Importinβ-1 or HEK293T_NL4.3 -Importinβ-1 viruses infected control and endogenous Importinβ-1 depleted TZM-bl cells, 8 hrs of infection. Quantification of capsid protein presence in the nucleus and cytoplasm of infected TZM-bl cells using ImageJ software. **(C)** Confocal microscopy for sub cellular localisation of capsid protein in the SupT1_NL4.3 +Importinβ-1 or SupT1_NL4.3 -Importinβ-1 viruses infected control and endogenous Importinβ-1 depleted TZM-bl cells, 6-7 hrs of infection. Quantification of capsid protein presence in the nucleus and cytoplasm of infected TZM-bl cells using ImageJ software. **(D)** Confocal microscopy for sub cellular localisation of capsid protein in the 1321N1_NL4.3 virus infected control and endogenous Importinβ-1 depleted TZM-bl cells, 6-7 hrs of infection. Quantification of capsid protein presence in the nucleus and cytoplasm of infected TZM-bl cells using ImageJ software. All experiments were done at least 3 times. capsid localisation quantification represents at least 10 cells from different fields. The significance is determined using an unpaired student's t-test. The p values are denoted as *** p <0.001; ** p <0.01, while non-significant values are denoted by n.s.

**Supplementary Figure S5.**

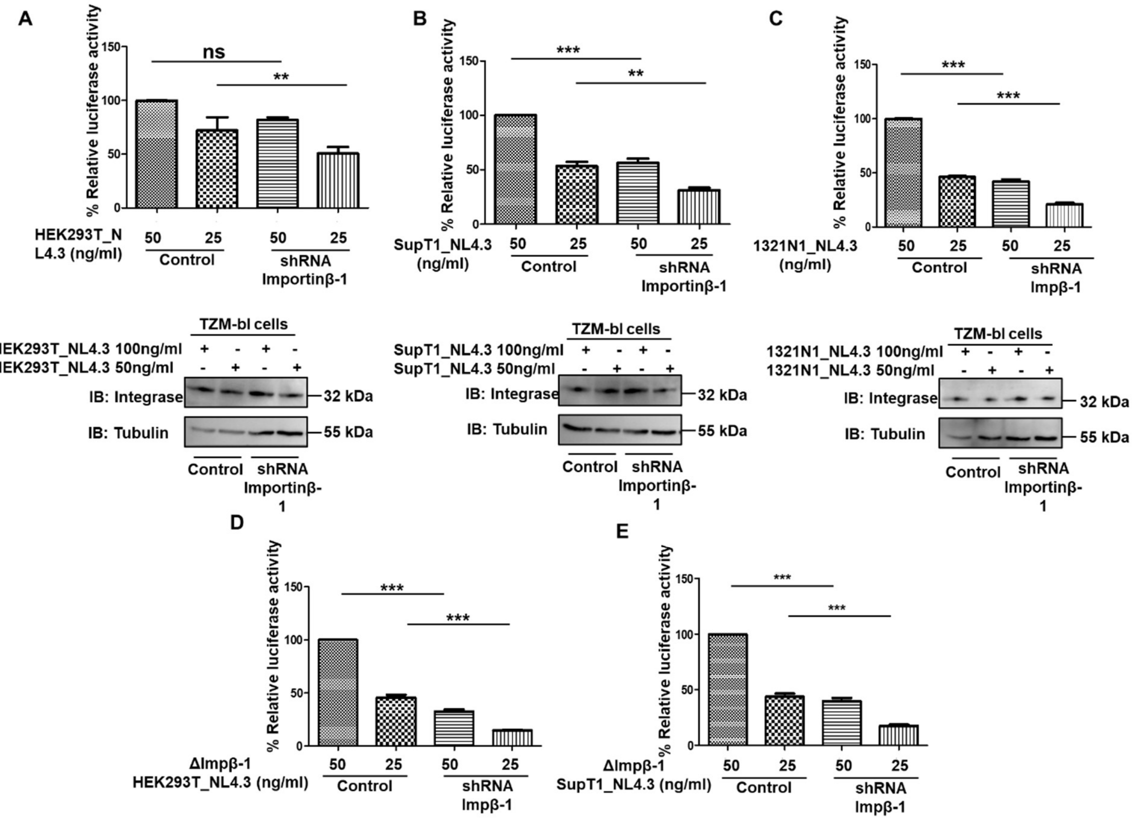


**Figure S5: Importinβ-1 presence in virion or cells is critical for HIV core entry. (A-C)** LTR-driven luciferase activity with HEK293T_NL4.3, SupT1_NL4.3, 1321N1_NL4.3 viruses respectively infected Importinβ-1 depleted and control TZM-bl cells. Besides, immunoblot representing the integrase protein in TZM-bl cells after infection with HEK293T_NL4.3, SupT1_NL4.3, 1321N1_NL4.3 viruses. **(D-E)** LTR-driven luciferase activity with ΔImportinβ-1 viruses of HEK293T_NL4.3, SupT1_NL4.3, viruses infected Importinβ-1 depleted and wild type TZM-bl cells, besides, immunoblot representing the integrase protein in TZM-b cells after infection with ΔImportinβ-1 of HEK293T_NL4.3, SupT1_NL4.3 viruses. All experiments were done at least 3 times. The significance is determined using an unpaired student's t-test. The p values are denoted as *** p <0.001; ** p <0.01, while non-significant values are denoted by n.s.

**Supplementary Figure S6. Importinβ-1 levels alter during HIV infection.**

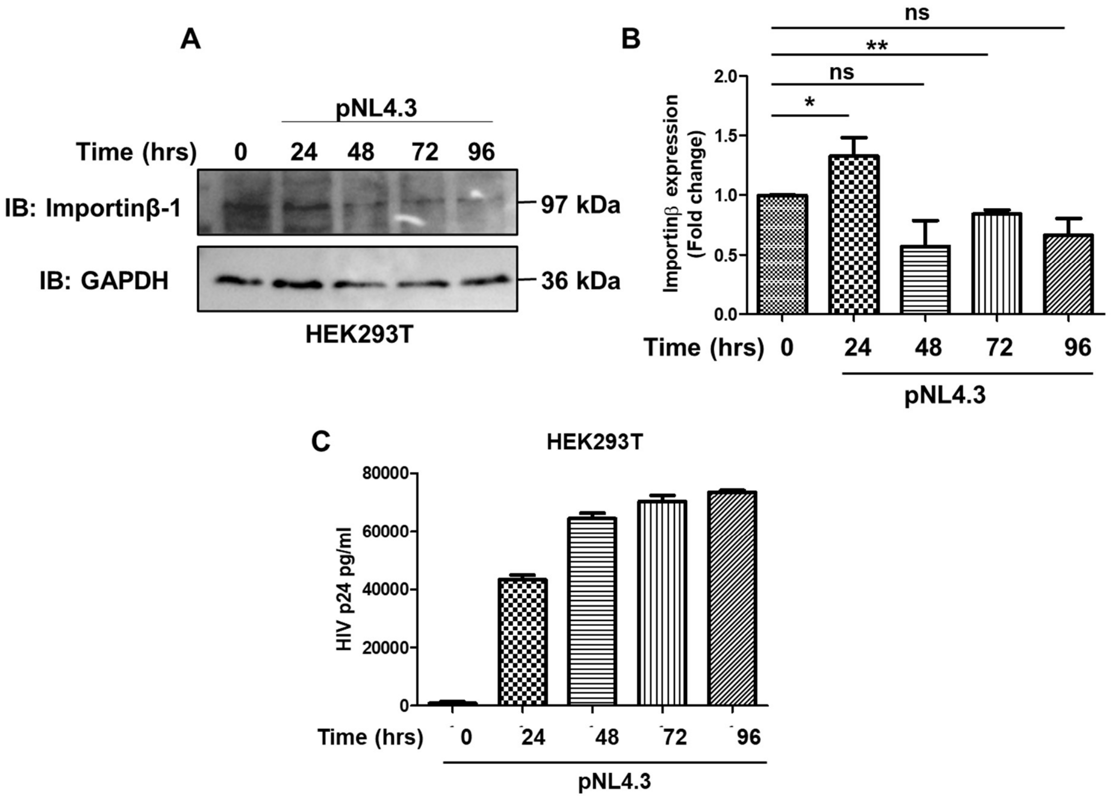


**Figure S6: Importinβ-1 levels alter during HIV infection. (A)** Representative immunoblot showing the levels of Importinβ-1 from 0hrs to 96hrs in HEK293T cells upon pNL4.3 transfection. GAPDH was used as a loading control. **(B)** Bar graph representing the quantification of Importinβ-1 expression in three independent experiments, including the experiment represented in A. **(C)** Levels of released virus p24 from 0 hrs to 96 hrs in pNL4.3 transfected HEK293T cells culture supernatant determined by p24 ELISA. All experiments were done at least 3 times. The significance is determined using an unpaired student's t-test. The p values are denoted as ** p <0.01; * p <0.05, while non-significant values are denoted by n.s.

**Supplementary Figure S7.**

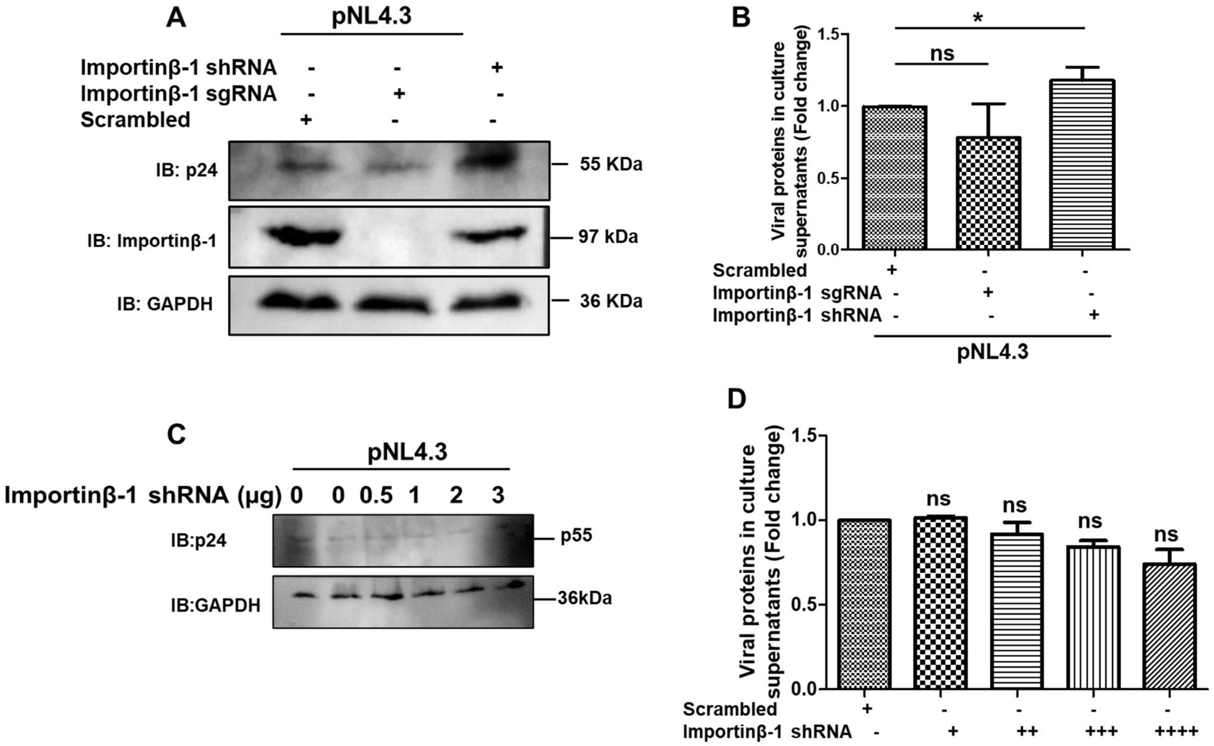


**Figure S7: Importinβ-1 knockdown enhances the cellular p55 levels and released viral titers.** **(A)** Representative immunoblot showing the p55 protein levels in the background of Importinβ-1 specific sgRNA, shRNA expressed HEK293T cells, GAPDH was used for the control. **(B)** Levels of released virus p24 in culture supernatant of pNL4.3 transfected HEK293T cells in the background of Importinβ-1 specific sgRNA, shRNA expression. The quantification was done by p24 ELISA. **(C)** Representative immunoblot showing the p55 protein levels in the background of gradually increasing the concentration of Importinβ-1 specific shRNA in HEK293T cells, GAPDH was used for the control. **(D)** Levels of released virus in culture supernatant of pNL4.3 transfected HEK293T cells upon gradually increasing the concentration of Importinβ-1 specific shRNA transduction. The quantification was done by p24 ELISA. All experiments were done at least 3 times. The significance is determined using an unpaired student's t-test. The p values are denoted as * $p < 0.05$, while non-significant values are denoted by n.s.

**Supplementary Figure S8.**

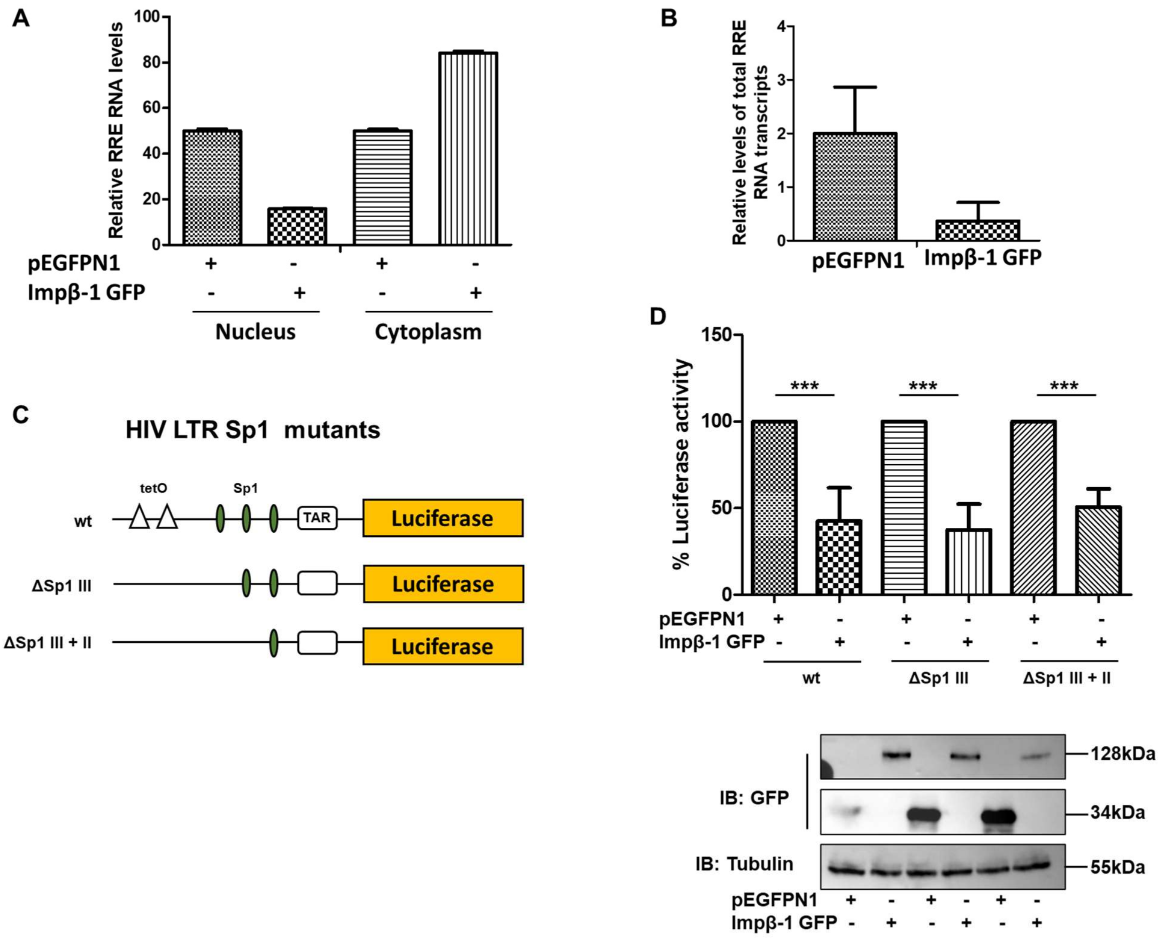


**Figure S8: Importinβ-1 inhibits the HIV transcription by regulating at the Sp1 site. (A)** qRT-PCR was performed with nuclear and cytoplasmic cDNA samples to check the RRE RNA levels in pNL4.3 transfected HEK293T cells transiently expressing the Importinβ-1 GFP. **(B)** Bar graph representing the total RRE RNA levels in pNL4.3 transfected HEK293T cells transiently expressing Importinβ-1 GFP. **(C)** Schematic representation of chimeric LTR Sp1 deletion mutants. **(D)** LTR driven luciferase activity in HEK293T cells upon transiently expressing Importinβ-1 GFP and chimeric SP1 mutants. Besides immunoblot showing the expression of Impβ-1 GFP and GFP. And Tubulin was used as a loading control. Experiments A, B were done twice and experiment D were done at least 3 times. The significance is determined using an unpaired student's t-test. The p values are denoted as *** $p < 0.001$, while non-significant values are denoted by n.s.

**Supplementary Figure S9.**

The *Insilco* binding analysis indicated that Sp1 region in LTR has a higher probability of binding to Importinβ-1 and we observed favourable Dock score and confidence scores for the Sp1-Importinβ-1 complex (Table S3). Additionally, Sp1 region occupied the larger cavity/pocket region of Importinβ-1 protein and formed many hydrogen bonds directly with the DNA bases, while others were established with the phosphate backbone. Further, the Sp1 and Importinβ-1 complex is stabilized by 17 hydrogen bonding interactions with Arg707, His309, Ser708, Ser750, Asp292, Glu289, Glu203, Glu296, Leu247, Asp288, Glu281, Asn285, Asp162, Leu247 and Tyr752 amino acid residues. Interestingly the amino acid residues majorly presented at the N-terminal and C-terminal domine of Importinβ-1. Overall, this Insilco analysis suggested Importinβ-1 has a DNA binding pocket and it can bind to the Sp1 region in the LTR (Figure S9A, B).

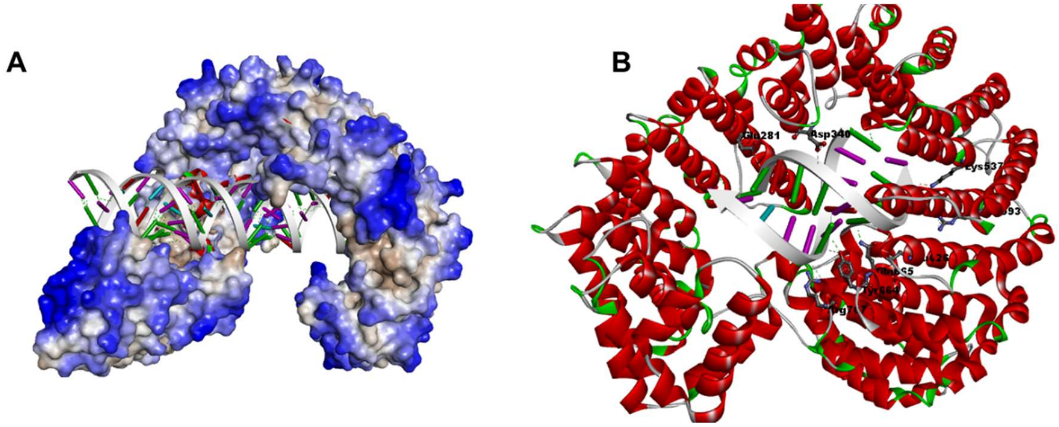


**Figure S9: Insilco binding prediction of Importinβ-1 and LTR regions. (A)** Insilco binding prediction of Importinβ-1 and Sp1 region of LTR by performing the docking studies. **(B)** Insilco binding prediction of Importinβ-1 and single Sp1 site of LTR by performing the docking studies.

**Table S3. Interaction studies of Importinβ-1 with various regions of LTR**

| DNA region | Confidence Score | Interacting residues |
|---|---|---|
| Sp1 region of LTR | DS= -342.25 Confidence Score=0.98 | B: LYS586:NZ - S: DT23:OP2, B: ARG800:NH2 - S: DA28:OP2, B: SER522: OG - A: DC15:OP2, B: THR578:OG1 - A: DC14:OP2, B: ARG585:NH1 - A: DG12:N3, B: TYR620: OH - A: DG12:OP2, B:SER841:CB - S:DG29:OP2, A:DC29:C4' - B:GLU239:OE1, 530B |

| | | |
|---|---|---|
| NFKB region of LTR | DS= -314.22 Confidence Score=0.92 | B: ARG585:NH1 - A: DC11:OP2, B: THR526:OG1 - S: DC5:OP2, B: TYR530: OH - A: DC14:OP2, B: ARG800:NH2 - A: DG6:O4', S: DG3:C4' - B: ASP440:OD2, S: DA4:C5' - B: ASN479:OD1, S: DA4:C5' - B: GLU482:OE1, S: DC5:C5' - B: THR526:OG1, S: DA18:C1' - B: ASN668:OD1, S: DT21:C1' - B: THR847:OG1, S: DT21:C5' - B: ASP808:OD1, S: DT22:C5' - B:ASP846:O, A: DA12:C5' - B: ARG585:O, A: DG13:C5' - B: GLU529:O |
| NRE region of LTR | DS= -220 Confidence Score=0.80 | A: LYS859:NZ - S: DG90:OP2, A:ARG863:NH1 - S:DG90:OP2, A: LYS867:NZ - S: DT91:OP2, A:GLN682:NE2 - S:DC101:OP2, A:ARG870:NH2 - S:DT91:O3’, A:LYS867:CE - S:DA92:OP2, S:DA102:C5' - A:ASP779:OD2, A:ARG863:NH2 - A:DT60:OP2 ,A: GLN491:NE2 - A: DT62:O4’, A: HIS597:CD2 - A: DT51:OP2, A:TRP864:CD1 - A:DC59:OP2, A: DT60:C5' - A: THR860:OG1 |
